# Exon-mediated activation of transcription starts

**DOI:** 10.1101/565184

**Authors:** Ana Fiszbein, Keegan S. Krick, Christopher B. Burge

**Author notes:** Correspondence/Lead Contact.

## Abstract

The transcription and processing of mammalian genes are intimately linked. Here we describe a phenomenon we call exon-mediated activation of transcription starts (EMATS), affecting thousands of genes, in which the splicing of internal exons impacts the spectrum of promoters used and expression level of the gene. We observed that evolutionary gain of new internal exons is associated with gain of new transcription start sites (TSS) nearby and increased gene expression. Inhibiting exon splicing reduced transcription from nearby promoters. Conversely, creation of new splice sites, enabling splicing of new exons, activated transcription from cryptic promoters. The strongest effects were associated with weak promoters located proximal and upstream of efficiently spliced exons. Together, our findings support a model in which splicing recruits transcription machinery locally to influence TSS choice, and identify exon gain, loss and regulatory change as major contributors to the evolution of alternative promoters and altered gene expression in mammals.

## Introduction

The processing of RNA transcripts from mammalian genes often occurs nearby in time and space to their synthesis, creating opportunities for functional connections between transcription and splicing (Barbosa-Morais et al., 2012; Merkin et al., 2012; Oesterreich et al., 2016; Osheim et al., 1985). Several links between splicing and transcription are known, and both transcription rate and chromatin structure can influence splicing outcomes in some cases (Bentley, 2014; Kornblihtt et al., 2013; Schor et al., 2013). However, more recent evidence suggests that splicing also feeds back on transcription (Braunschweig et al., 2013). Adding an intron to an otherwise intron-less gene often boosts gene expression in plants, animals, and fungi; the mechanisms are not fully understood but impacts on transcription, nuclear export, mRNA stability, and/or translation have been noted (Furger et al., 2002; Shaul, 2017). Splicing can impact transcription elongation rates (Fong and Zhou, 2001), and in yeast the presence of an intron can generate a transcriptional checkpoint that is associated with pre-spliceosome formation (Chathoth et al., 2014). Furthermore, recruitment of the spliceosome complex can stimulate transcription initiation by enhancing preinitiation complex assembly (Damgaard et al., 2008), and inhibition of splicing can reduce levels of histone 3 lysine 4 trimethyl (H3K4me3), a chromatin mark associated with active transcription (Bieberstein et al., 2012).

Several components of the splicing machinery associate with RNA polymerase II (RNAPII) and other transcription machinery (Das et al., 2007; Emili et al., 2002; Kameoka et al., 2004; Morris and Greenleaf, 2000; Mortillaro et al., 1996; Neugebauer and Roth, 1997; Vincent et al., 1996). The U1 and U2 small nuclear ribonucleoprotein particles (snRNPs) associate with general transcription factors (GTFs) GTF2H (Kwek et al., 2002), GTF2F (Kameoka et al., 2004), and the carboxy-terminal domain (CTD) of RNAPII (Emili et al., 2002; Morris and Greenleaf, 2000). In addition to its role in splicing, U1 snRNP acts as a general repressor of proximal downstream premature cleavage and polyadenylation (PCPA) sites (Gunderson et al., 1998; Kaida et al., 2010). The relative depletion of U1 snRNP binding sites upstream in the antisense orientation from promoters (relative to their presence in the downstream sense direction) contributes to frequent termination of antisense transcripts at PCPA sites, resulting in short unstable transcripts (Almada et al., 2013).

Alternative start and termination sites drive a substantial portion of transcript isoform differences between human tissues (Reyes and Huber, 2018). Recent analyses of full-length mRNAs suggests that transcription starts and splicing may be coordinated (Anvar et al., 2018). However, whether exon splicing commonly impacts transcription start site (TSS) location and activity remains unknown. Here we describe a phenomenon we call “exon-mediated activation of transcription starts” (EMATS) in which the splicing of internal exons, especially those near gene 5’ ends, alters gene expression by influencing which TSSs are used. Our results demonstrate that exon splicing activates transcription initiation locally in thousands of mammalian genes, including many involved in brain development. Our findings also indicate that activation or repression of gene expression for research or therapeutic purposes may commonly be achievable by manipulation of splicing.

## Results

### Increased exon splicing is associated with increased gene expression and alternative TSS usage

We used a comparative approach to explore potential connections between splicing and TSS usage, examining transcript patterns in orthologous genes of mouse and rat that differed by the presence/absence of an internal exon (i.e. a non-terminal exon flanked by introns). Previously, we identified over one thousand such exons that were unique to the mouse transcriptome and not detected in RNA-seq data from diverse organs/tissues of other mammals including rat, macaque, cow, etc., and therefore likely arose recently in the mouse lineage. We also identified a similar number of exons that are unique to the rat. Most such evolutionarily new exons are located in 5’ untranslated regions (UTRs) and are spliced in an alternative and tissue-specific fashion (Merkin et al., 2015). Comparing closely related species, we have observed that genes with evolutionarily new internal exons tend to have increased gene expression, but only in those tissues where the new exons are included in mRNAs (Figure 1A, S1A and Table S1) (Merkin et al., 2015). This trend was stronger for exons that were efficiently spliced – assessed by “percent spliced in” (PSI or ψ) values > 0.95, indicating that more than 95% of mRNAs from the gene include the exon (Figure 1B) – suggesting an association between the extent of exon splicing and level of gene expression.

**Figure 1.**
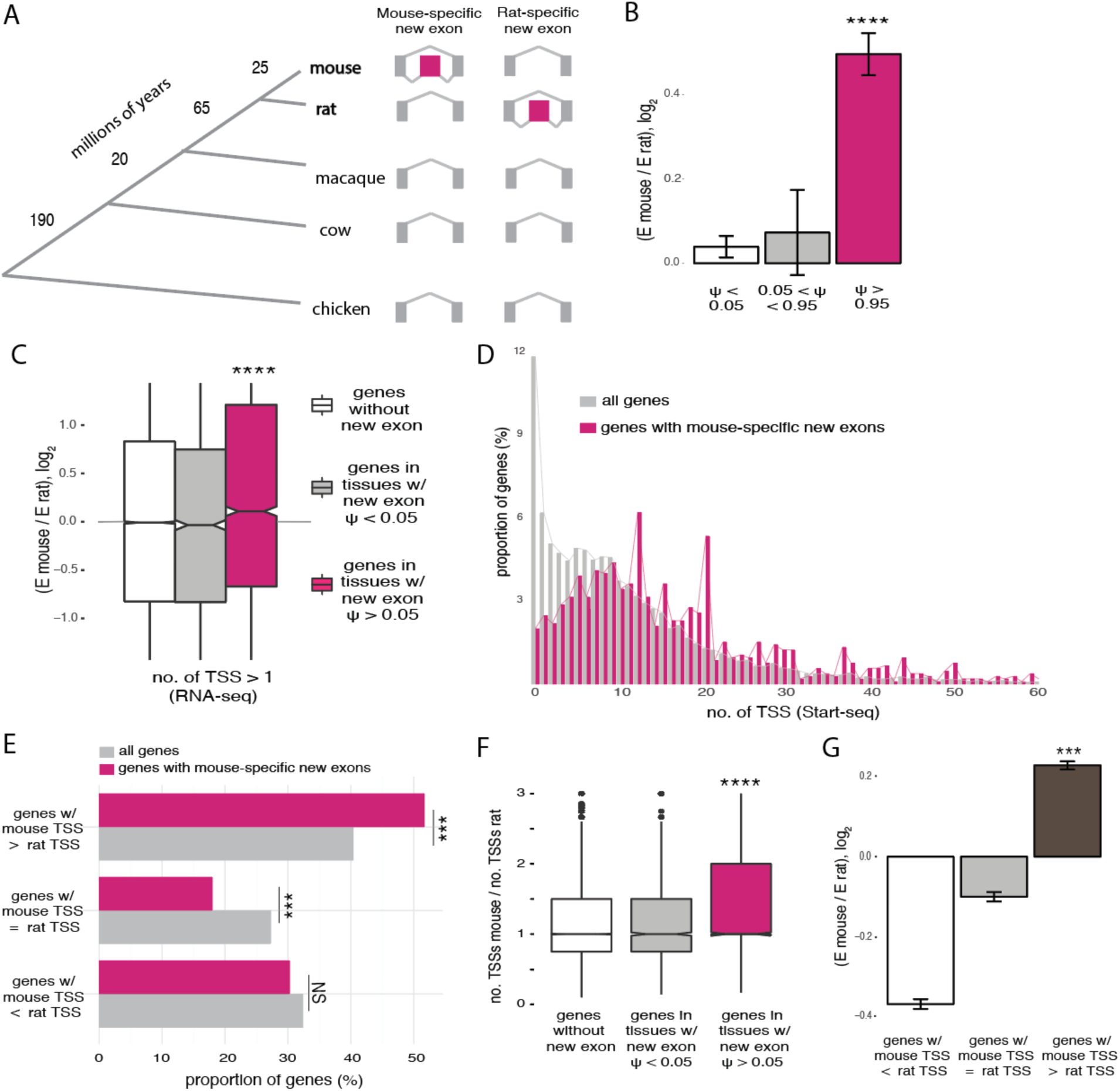
Splicing is associated with increased gene expression and usage of TSS. A, Phylogenetic tree representing the main species used for dating evolutionarily new exons and approximate branch lengths in millions of years. The patterns of inclusion/exclusion used to infer mouse-specific new exons (*n* = 1089) and rat-specific new exons (*n* = 1517) are shown. B, Fold change in gene expression between mouse and rat in 9 organs (brain, heart, colon, kidney, liver, lung, skeletal muscle, spleen and testes) for genes with mouse-specific new exons, binned by ψ value of the new exon in each tissue. **** p < 0.0001 by one-way ANOVA, Tukey post hoc test. C, Fold change in gene expression between corresponding tissues of mouse and rat in genes with multiple TSSs in mouse (no. of TSS > 1) for mouse control genes with no new exons (white), genes with mouse-specific new exons in tissues where inclusion of the new exon is not detected, PSI < 0.05 (grey), and genes with new mouse-specific exons in tissues were the exon is included, PSI > 0.05 (pink). *** p < 0.001 by one-way ANOVA, Tukey post hoc test. D, Distribution of the number of TSSs per gene using Start-seq data from murine macrophages for all genes expressed in mouse and genes with mouse-specific new exons. TSS peaks located within 50 bp apart were merged. Genes with mouse-specific new exons have increased numbers of TSSs (p < 2.2e^−16^ by Kolmogorov-Smirnov test). E, Proportion of genes that gained TSSs in mouse (mouse TSS > rat TSS), genes that lost TSSs in mouse (mouse TSS < rat TSS) and genes with same number of TSSs in both species (mouse TSS = rat TSS) for all genes expressed in both species and genes with mouse-specific new exons. *** p < 0.001 by one-way ANOVA, Tukey post hoc test. F, Fold change in the number of TSSs used per gene between mouse and rat for 9 tissues, for mouse genes grouped as in (C). G, Fold change in gene expression between mouse and rat for genes grouped as in (E). See also Figure S1.

Grouping genes by their promoter structure, we found a positive association between inclusion of new exons and gene expression for genes with multiple TSSs while this association was not observed for genes with only one TSS (Figure 1C). Furthermore, our RNA-seq data (from (Merkin et al., 2012)) showed that genes with mouse-specific new exons were far more likely to have multiple TSSs compared to all expressed genes in mouse (Figure S1B and S1C). We confirmed that genes with new mouse-specific exons are more likely to have multiple TSSs, using other methods to define TSS locations, including H3K4me3 ChIP-seq peaks (Yu et al., 2015) and data from high-resolution sequencing of polymerase-associated RNA (Start-seq) (Scruggs et al., 2015) (Figure 1D, S1D, Table S2). Genes with rat-specific new exons also had more TSSs per gene than rat genes overall (Figure S1E). Furthermore, genes that gained new species-specific exons were more likely to have gained TSSs in the same species, suggesting that the evolutionary gain of an internal exon is connected to evolutionary gain of TSSs in a locus (Figure 1E and S1F).

To investigate this connection further, we examined the usage of new exons and TSSs used by a gene in different tissues. We observed that genes containing mouse-specific exons used more distinct TSSs than their rat orthologs (Figure S1G), and that this association was specific to mouse tissues where the new exon was included with PSI > 0.05 (Figure 1F), showing a connection between splicing and TSS use in different mammalian organs. We also observed higher PSI values for new exons in genes with multiple alternative TSSs relative to genes with a single TSS (Figure 1H). Furthermore, the increased gene expression levels in mouse relative to rat in genes with new mouse exons was restricted to those genes that gained TSSs in mouse (Figure 1G and S1I). Together, these observations indicate that the usage of new TSSs and the splicing of new internal exons tend to occur in the same genes, tissues, and species, suggesting an intimate connection between splicing, increased gene expression and activation of new TSSs.

### TSSs arise proximal and upstream of new exons

We observed a positional effect in which the increase in TSS counts per gene was associated predominantly with new exons located in 5’ UTRs rather than those in 3’ UTRs or coding regions (Figure S2A). We examined the distribution of the locations of all mouse TSSs relative to the locations of mouse-specific new exons (Figure S2B), and compared it to the distribution of rat TSSs relative to sites homologous to the mouse-specific exons. This comparison showed an enrichment of TSSs in mouse within one or two kilobases (kb) upstream of new exons (Figure 2A, inset and Figure S2C). Thus, evolutionary gain of new internal exons was specifically associated with gain of proximal, upstream TSSs.

**Figure 2.**
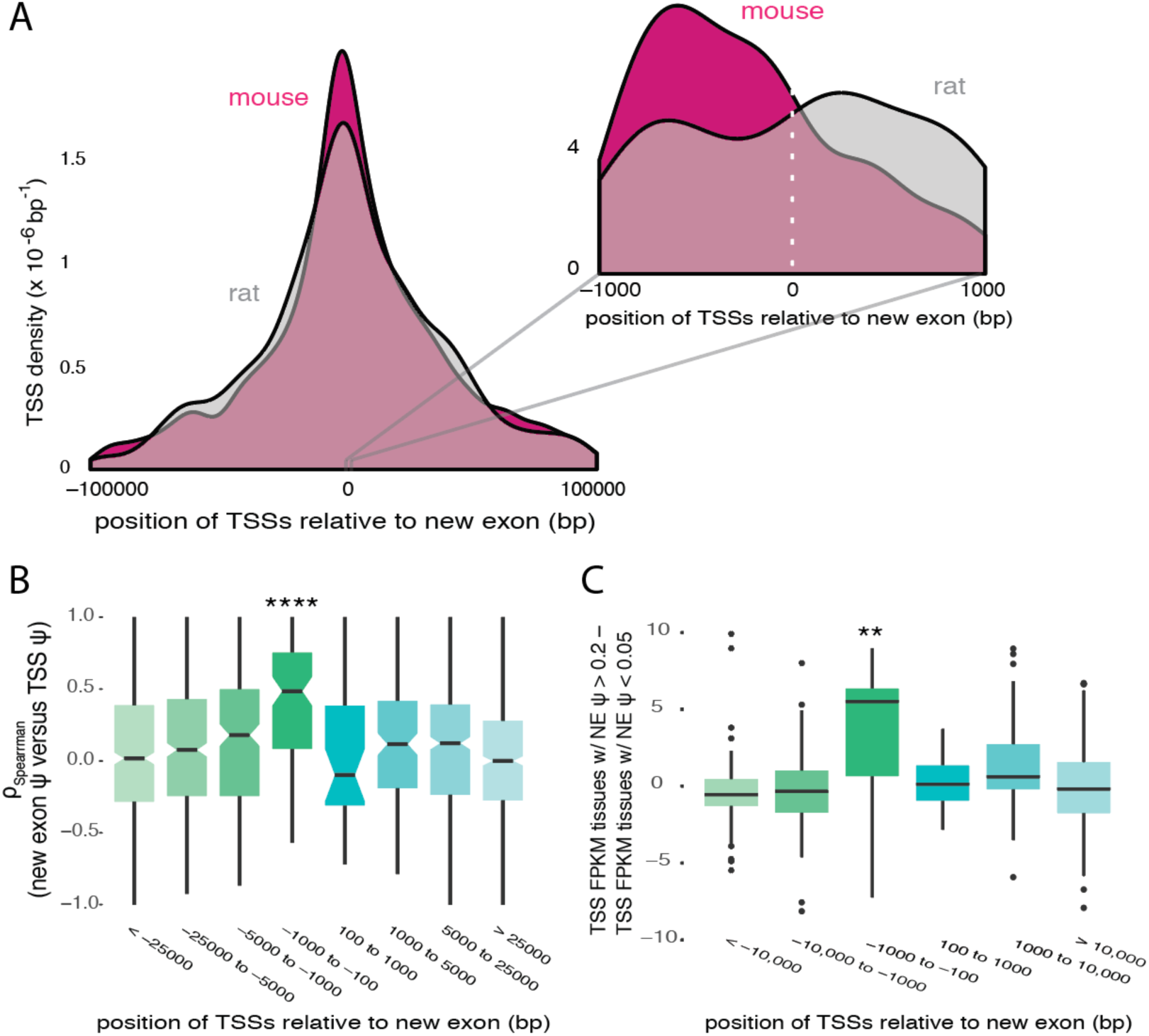
TSSs arise proximal and upstream of new exons. A, Histogram of TSS locations in mouse (pink) and rat (grey) in all 9 tissues for genes with mouse-specific new exons, centered on start of mouse new exon or homologous genomic position in rat. Inset zooms in on locations within 1 kb of new exon. Distributions were smoothed with kernel density estimation by ggplot2 with default parameters. B, Spearman correlations between TSS PSI and new exon PSI across mouse tissues, for TSSs binned by position relative to mouse-specific exon. C, Difference in expression (in units of fragments per kilobase of exon per million mapped reads, FPKM) in mouse tissues for transcripts including TSSs in tissues where new exon is moderately or highly included (PSI > 0.2) versus tissues where new exon is excluded (PSI < 0.05), grouped by TSS location relative to new exon. See also Figure S2.

We then asked about the relationship between splicing levels and usage of alternative TSS within the same gene. Considering relative TSS use (“TSS PSI”, representing the fraction of transcripts from a gene that derive from a given TSS) we found that use of the most proximal upstream TSS (designated TSS –1) was positively correlated with new exon inclusion, especially for TSSs located within about 1 kb upstream of the new exon (Figure 2B and S2D). Furthermore, absolute expression of transcripts from nearby TSSs increased specifically in tissues where new exons were included at moderate or high levels (Figure 2C). These observations suggest a positive influence of splicing on nearby transcription (or possibly vice versa).

### Manipulation of exon splicing impacts upstream transcription initiation

To directly test whether splicing impacts nearby transcription, we chose two candidate mouse genes, *Gper1* (G protein-coupled estrogen receptor 1), and *Tsku* (Tsukushi, small leucine rich proteoglycan). These genes both have widespread, moderate expression and contain a mouse-specific 5’ UTR internal exon whose splicing is positively correlated with the expression of the gene across mouse tissues (Spearman ρ = 0.64 and 0.57, respectively; Figure 3A and 3B left panels). When cultured mouse fibroblasts were treated with morpholino antisense oligonucleotides (MO) targeting splice sites of the new exons, exon inclusion decreased by about 4-fold in both *Gper1* (Figure 3A) and *Tsku* (Figure 3B). Moreover, gene expression levels of these two genes were depressed to a similar extent (Figure S3A), consistent with a positive effect of exon inclusion on gene expression. We observed similar levels of repression when assaying metabolically labeled nascent RNA (Figure S3A) as with total mRNA, indicating that the effect is primarily at the level of transcription rather than mRNA stability (Figure 3A and 3B). These observations support the idea that exon splicing positively impacts nearby transcription.

**Figure 3.**
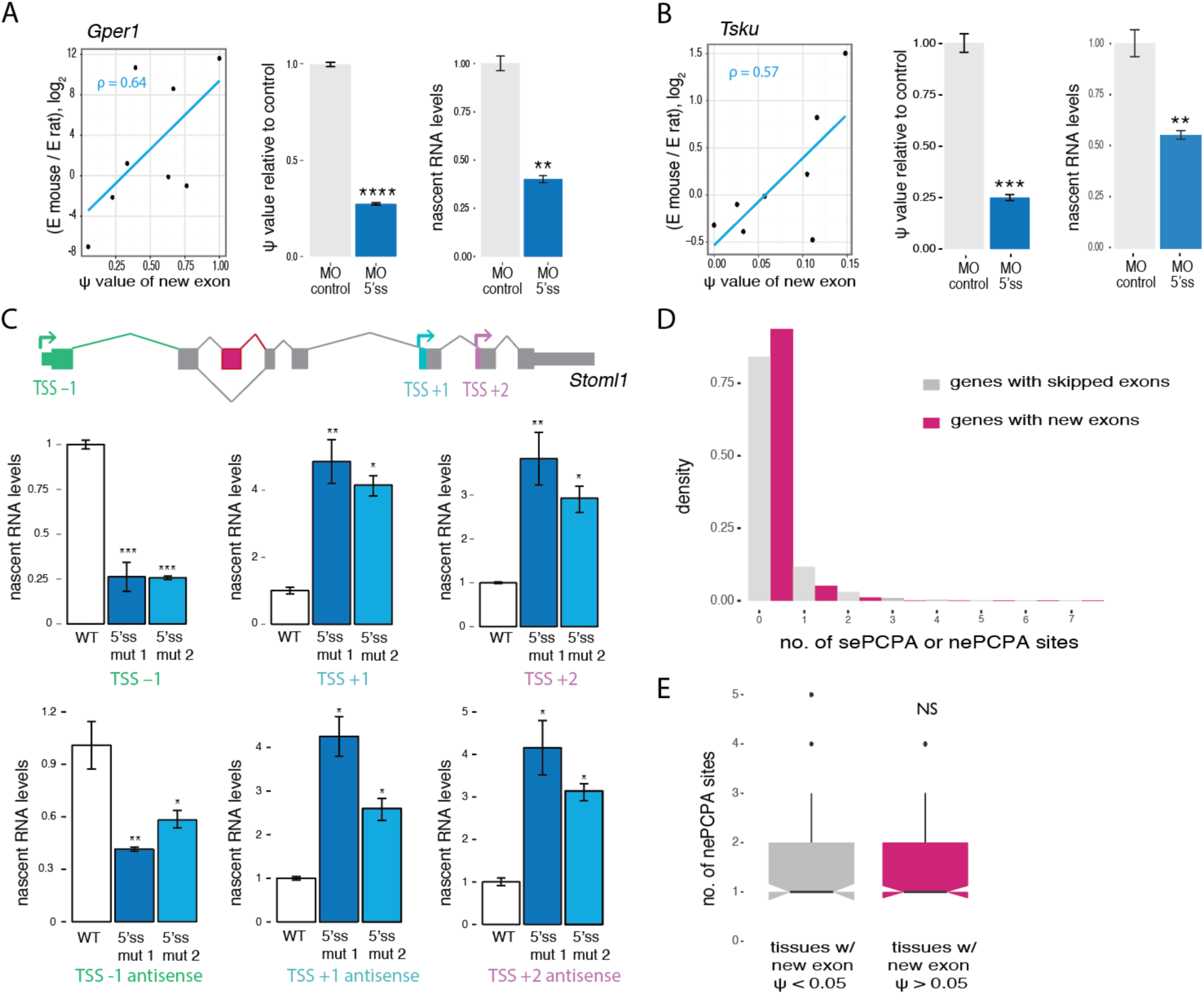
Manipulation of exon splicing impacts upstream transcription initiation. A, (Left) Relationship between fold change in gene expression between mouse and rat and new exon PSI value across 9 tissues for *Gper1* gene. (Right) qRT-PCR analysis of fold change in new exon PSI value (middle) and gene expression (right) in nascent RNA metabolically labeled for 10 minutes with 5-ethynyl uridine, following treatment of NIH3T3 cells with MO targeting new exon 5’ splice site relative to control treatment. Mean ± SEM of displayed distributions, *n* = 3 biological replicates. ** p < 0.01, *** p < 0.001,****p < 0.0001 by one-way ANOVA, Tukey post hoc test. B, As in (A) for mouse *Tsku* gene. C, Fold change in nascent sense (top) and antisense (bottom) RNA levels of *Stoml1* in CAD cells measured by qRT-PCR of RNA metabolically labeled for 10 minutes with 5-ethynyl uridine and normalized using housekeeping genes *Gapdh, Hprt* and *Hspcb*. Wild type cells in white and CRISPR-Cas cells with mutations in the 5’ splice site of the new exon in blue. Mean ± SEM of displayed distributions, *n* = 3 independent experiments. A schematic diagram of *Stoml1* exon-intron organization is shown at top. D, Distribution of the number of polyadenylation sites used per gene located 2 kb upstream/downstream of a control set of mouse genes with skipped exons (grey, sePCPA) and genes with mouse-specific new exons (pink, nePCPA). sePCPA and nePCPA are defined in Methods. E, Distribution of the number of polyadenylation sites used 2 kb upstream/downstream of new exons per gene in tissues where new exon is excluded (PSI < 0.05, grey) or included (PSI > 0.05, pink), for genes with new exons and at least one nePCPA. Distributions are not significantly different by Kolmogorov-Smirnov test. See also Figure S3.

We next sought to confirm the directionality of this effect and to ask how splicing of new exons impacts the use of different TSSs. We chose for analysis the mouse *Stoml1* (Stomatin Like 1) gene, because it has three alternative TSSs as well as a new exon, all of which are used in mouse fibroblasts (Figure S3C). Using CRISPR/Cas9 mutagenesis to generate cell lines with mutations abolishing the inclusion of a new exon we observed that the three alternative TSSs of the gene responded differently to inhibition of splicing of the new exon. The upstream –1 TSS was down-regulated by 4-fold, while downstream +1 and +2 TSSs were up-regulated to a similar extent in the mutant cell lines as measured by qPCR of nascent RNA (Figure 3C). Effects on antisense transcription in these mutant cell lines mirrored those observed for sense transcription (Figure 3C), suggesting that inclusion of the new exon enhances transcription from the upstream promoter in both directions. This pattern is distinct from a report of intron-mediated enhancement in which sense-oriented introns specifically inhibited antisense transcription (Agarwal and Ansari, 2016), but is consistent with reported impacts on transcription initiation resulting from changes in the position of an intron in a reporter gene (Gallegos and Rose, 2017). The increase in downstream promoter activity was not observed in other genes studied and may reflect some sort of locus-specific (e.g., homeostatic) regulation of *Stoml1* expression. Levels of H3K4me3 and RNAPII decreased in the upstream TSS and increased in the downstream TSSs in the mutant cell lines, consistent with the observed effects on nascent transcript production (Figure S3C).

Premature cleavage and polyadenylation can produce truncated, unstable transcripts, but can be inhibited by binding of U1 snRNP near of a PCPA site (Gunderson et al., 1998; Kaida et al., 2010). If the observations above reflected effects of U1 snRNP or other splicing machinery on PCPA rather than on transcription, this would require the presence of new exon-proximal PCPA (“nePCPA”) sites in affected genes. Using available polyA-seq data from five mouse tissues (Methods), we observed that only 8.6% of genes with new exons had evidence of a nePCPA site, slightly lower than in a control set of genes (Figure 3D). And for the subset of genes that contain nePCPA site(s), we did not observe differences in usage of the site between tissues where the new exon was spliced in and those where it was spliced out (Figure 3E and S3D). Furthermore, we saw no relationship between the number of nePCPA sites and gene expression changes between mouse and rat (Figure S3E). Thus we found no evidence that effects on PCPA contribute significantly to EMATS. Since we observed similar effects on nascent RNA (in both sense and antisense orientations) as on total mRNA levels, our results imply that EMATS impacts transcription initiation rather than later steps.

### Creation of a new splice site activates the use of a cryptic promoter nearby

We next sought to explore how splicing might affect use of different upstream TSSs. In the *Tsku* gene, the mouse-specific TSS in position –1 is located within 1 kb upstream of the mouse-specific exon, while the conserved TSS –2 is located further upstream. Analysis by 5’ RACE showed that both TSSs are normally used at similar levels in mouse fibroblasts. However, inhibiting splicing of the new exon by MO resulted in lower use of TSS –1 (Figure 4A and S4A). The strong down-regulation of transcription from TSS –1 observed by 5’ RACE was confirmed by qRT-PCR of nascent RNA, in both sense and antisense orientations (Figure S4B). This shift was accompanied by a 3-fold decrease in H3K4me3 levels near TSS –1 in MO-treated cells (Figure 4B). However, levels of H3K4me3 near TSS –2 were unchanged, confirming that TSS – 2 transcription is not affected (Figure 4B). In cells treated with MOs, levels of GTF2F1 and RNAPII decreased by almost 3-fold near TSS –1 but were unchanged near TSS –2 (Figure 4C and S4C). These observations suggest that splicing of the new exon contributes to recruitment of core transcription machinery to the proximal TSS –1. Moreover, the loss of signal for GTF2F1 and RNAPII near the new exon following MO treatment suggests that inclusion of the new exon is associated with recruitment of transcription factors and higher levels of RNAPII, consistent with functional interactions between GTFs and splicing machinery observed previously (Damgaard et al., 2008; Das et al., 2007). These observations confirm that splicing of new exons can regulate the usage of alternative TSSs, with predominant effects on proximal upstream promoters, consistent with the correlations observed in Figure 2.

**Figure 4.**
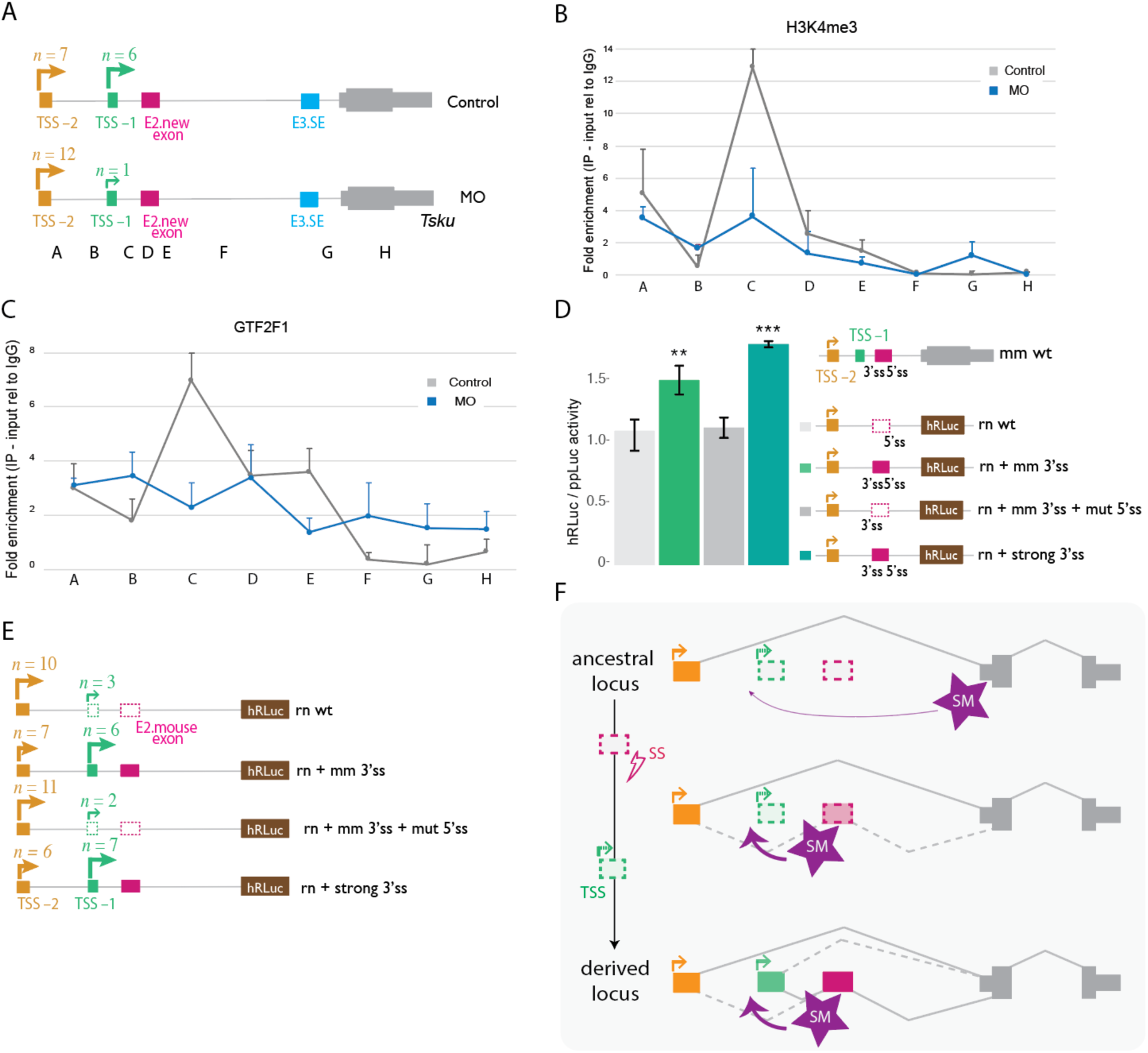
Creation of a new splice site activates the use of a cryptic promoter nearby. A, Schematic of 5’ RACE products and number of clones obtained for each TSS in control NIH3T3 cells and cells transfected with MO targeting the 3’ and 5’ splice sites of the new exon in *Tsku, n* = 2 biological replicates. B,C, ChIP-PCR analysis of H3K4me3 (B) and GTF2F1 (B) in *Tsku* gene in NIH3T3 cells for regions indicated in (A). Mean ± SD of two independent immunoprecipitations normalized to input and mean value for control IgG antibody are shown. Data shown for control cells (grey) and cells treated with MOs targeting the 3’ and 5’ splice sites of the new exon (blue). D, Luciferase activity in HeLa cells transfected with the hybrid constructs of the *Tsku* gene (right). Promoter activities of the corresponding constructs (corrected for transfection efficiency) are presented as fold increase of *Renilla* luciferase activity relative to firefly luciferase activity (encoded on the same plasmid). Mean ± SD for *n* = 3 independent experiments. E, Schematic of 5′ RACE analysis showing TSS usage and number of clones obtained in NIH3T3 mouse cells transfected with plasmids expressing the corresponding rat *Tsku* mutants. F, Model in which creation of a splice site during evolution triggers inclusion of a new internal exon which activates use of an upstream cryptic TSS. In the model, exon recognition by the splicing machinery (SM) in transcripts from the distal promoter activates TSS(s) located proximal and upstream of the exon. Transcripts initiating from the proximal promoter also include the exon, further boosting activity of this promoter. See also Figure S4.

To dissect the impacts of individual splice sites and splicing levels, we created an exon corresponding to the mouse-specific new exon in the rat *Tsku* gene and assessed effects on transcription. In the rat *Tsku* locus, transcripts are predominantly transcribed from the distal TSS –2. However, the regions homologous to TSS –1 and the mouse-specific new exon have high sequence identity with the mouse genome: both 5’ splice sites are present in rat, but no YAG is present in rat near the location of the mouse 3’ splice site, likely preventing splicing (Figure S4D). To introduce the desired mutations, we cloned the 5’ end of the rat *Tsku* gene upstream of the coding sequence of *Renilla* luciferase and recreated the 3’ splice site that is present in the mouse genome (rn + mm 3’ss), as well as a stronger 3’ splice site (rn + strong 3’ss), while either maintaining or mutating the native rat 5’ splice site sequence (mm 3’ss + mut 5’ss). Strikingly, the creation of a 3’ splice site promoted the inclusion of an exon analogous to that observed in mouse in constructs with an intact 5’ splice site (Figure S4E), indicating that this mutation is sufficient to create a new exon in the rat gene. In the presence of both 3’ and 5’ splice sites, but not when either splice site was absent, total gene expression levels increased, as measured by luciferase activity (Figure 4D). By 5’ RACE analysis, TSS –1 is used at basal levels in the minigene. However, the mouse-specific exon activates the usage of TSS –1 by 3-fold in the presence of a 5’ splice site, demonstrating that the effect on TSS usage depends on splicing of the mouse-specific exon rather than merely the presence of a 3’ splice site sequence (Figure 4E and S4F).

In some examples studied previously, species-specific alternative splicing alters protein function (Gracheva et al., 2011; Gueroussov et al., 2015). Our observations support the existence of a distinct evolutionary pathway in which, following a mutation that generates a new internal exon, splicing of the new exon in transcripts from a distal upstream promoter activates a cryptic proximal upstream promoter; and transcripts from the new promoter also include the exon, further activating the new promoter in a sort of positive feedback loop. The resulting new TSS produces novel transcript isoforms and higher gene expression in tissues where the upstream promoter is active and the exon is included (Figure 4F). Conversely, loss of an internal exon may result in loss of a weak upstream promoter that is dependent on splicing of the exon.

### Efficiently spliced exons activate use of weak proximal TSSs

To investigate the genomic scope of the relationship between splicing and alternative TSS usage, we asked whether the inclusion of alternative skipped exons (SE) in general – not just those that evolved recently – can influence start site selection. We identified 49,488 SEs in mouse RNA-seq data, distributed across 13,491 genes using conservative criteria (Table S3). Analyzing unique SEs with TSS-exon distances matching those of new exons, we observed no significant association between SE inclusion and use of proximal upstream TSSs overall (Figure 5A). In addition, we observed a symmetrical distribution of TSSs around the locations of SEs, distinct from the upstream-biased distribution seen relative to new exons (Figure 5B). These differences suggest that genes with new exons have distinct properties that favor linkage of splicing with transcription.

**Figure 5.**
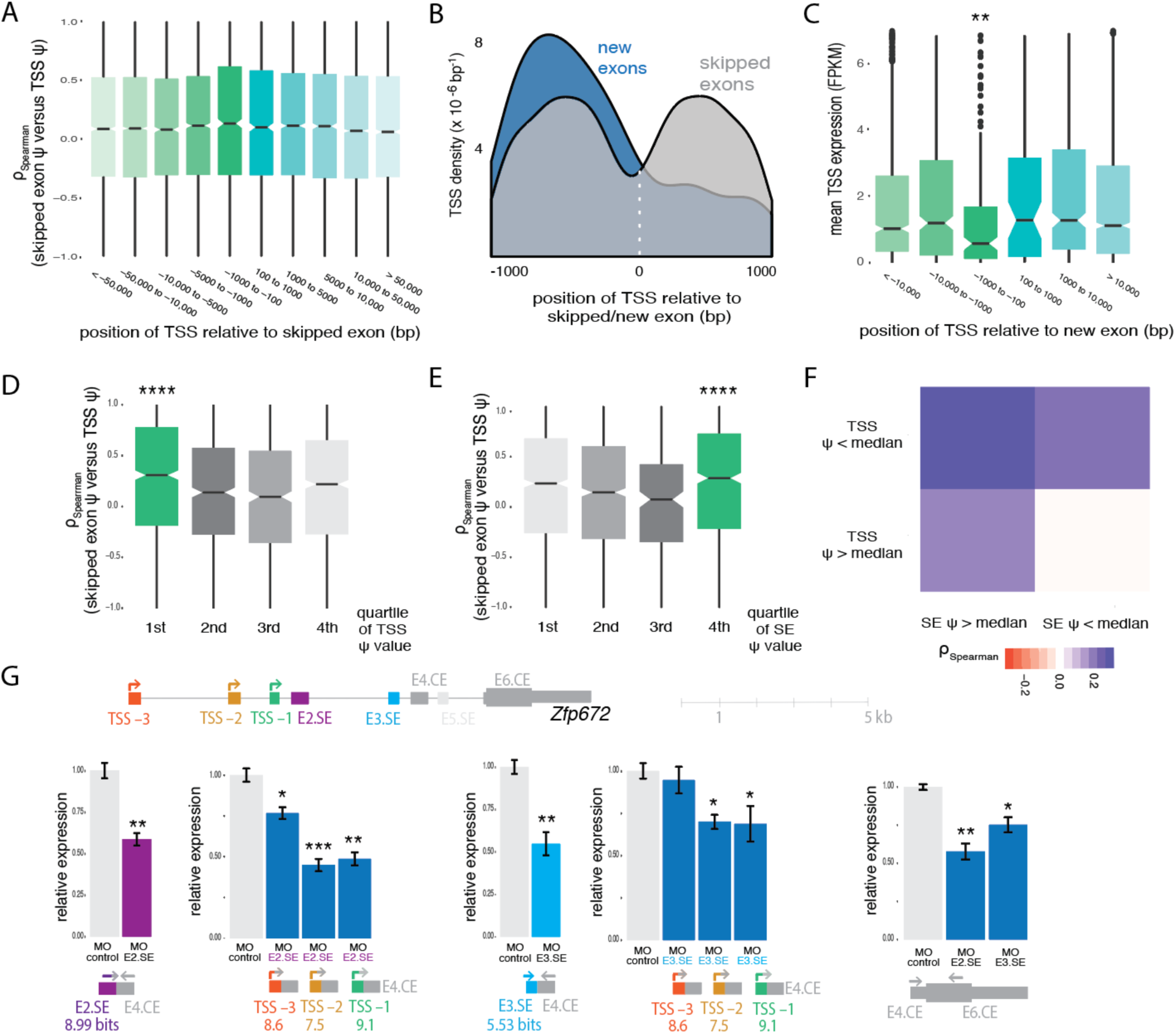
Efficiently spliced exons activate usage of weak proximal TSSs. A, Spearman correlations between TSS PSI (*n* = 49,911) and skipped exon PSE (SE, *n* = 13,491) in the same gene across mouse tissues for all expressed TSSs in genes with SEs, binned by genomic position relative to the SE. B, Comparison of distributions of TSS positions in 9 tissues for genes with mouse-specific new exons (pink) and genes with SEs in mouse (grey). Position 0 is set to the start coordinate of the new exon/skipped exon. Distributions were smoothed with Kernel density estimation. C, Expression of alternative first exons (AFE) for all TSSs in genes with mouse-specific new exons in tissues where the new exon is included (PSI > 0.05), binned by position relative to the new exon. D, Spearman correlation between TSS PSI and SE PSI in the same gene across mouse tissues for TSSs within 1kb upstream of the SE, binned by quartiles of mean TSS PSI. E, Same as (D) but binned by quartiles of mean SE PSI. F, Heat map showing the median Spearman correlation between TSS PSI and SE PSI in the same gene across mouse tissues for SEs with at least one TSS located upstream, in four groups, according to whether the mean TSS PSI (across tissues) and the mean SE PSI were greater than or less than the corresponding median values (across all TSSs and SEs analyzed). G, Exon-intron organization of mouse *Zfp672* gene. qRT-PCR analysis of expression of *Zfp672* in NIH3T3 cells normalized to expression of housekeeping genes *Hprt* and *Hspcb*. Data for control cells and cells treated with MO targeting the indicated splice sites (E4.CE and E6.CE). E5.SE is not included in NIH3T3 cells. Inclusion levels of the skipped exons, as well as levels of exon-excluding transcripts from the alternative TSSs (TSS –3, TSS –2, TSS –1) and total gene expression are shown. Scores of 5’ splice sites of skipped exons and first exons are listed in bits. Mean ± SEM of displayed distributions for *n* = 3 independent experiments. See also Figure S5.

Examining other features of loci with new exons, we observed that, although new exons tend to have lower PSI values than SEs overall (Figure S5A), those new exons with proximal upstream TSSs tended to have higher PSI values and stronger 5’ splice sites (Figure S5B). Furthermore, although the distribution of TSS PSI values was similar in genes with new exons and genes with SEs generally (Figure S5C), those TSSs located proximal and upstream of new exons had lower average expression levels across tissues than TSSs in other locations (Figure 5C). These observations suggested that the link between splicing and TSS usage is most pronounced when the promoter is intrinsically weak and splicing activity is high. Consistently, previous studies have observed stronger intron-mediated enhancement in the presence of weaker promoters (Callis et al., 1987). To test this idea, we grouped SEs and their most proximal and upstream TSS into four bins from weak to strong on the basis of the TSS PSI value, and separately for the SE PSI value, and analyzed the correlation between TSS PSI and SE PSI separately for each bin. Notably, we observed that TSS usage was most highly correlated with exon inclusion for the lowest quartile of TSS PSI values (Figure 5D, S5D and S5E) and for the highest quartile of SE PSI (Figure 5E and S5F). Thus, we found evidence that the EMATS observed for new exons occurs for a subset of general SEs. Robust effects were observed when a weak promoter is located upstream of a highly included SE, which occurred in 3,833 mouse genes, with the strongest effects seen for proximal weak promoters, which occurred in 1777 mouse genes (Figure 5F and Table S3). In humans, we identified 3548 genes with EMATS organization and 1098 genes with EMATS structure in which the weak promoter is also proximal to the SE (within 2 kb). Considering constitutive exons the number of identified genes increases by 3-fold.

To further investigate the distance dependence of splicing effects on TSS use, we analyzed changes in TSS usage when inhibiting the inclusion of a SE in the mouse *Tsku* locus located more than 6 kb downstream of the TSSs. Perturbations of the splicing of this exon caused no detectable changes in TSS usage (Figure S5G), consistent with a requirement for proximity of spliced exon to TSS for EMATS activity. Considering another mouse gene, *Zfp672* (Zinc Finger Protein 672) – chosen because it contained multiple TSSs and SEs expressed in mouse fibroblasts – we observed that inhibition of the stronger upstream SE in the locus affected the usage of TSSs more dramatically than inhibition of the weaker downstream SE in the same gene (Figure 5G). A weaker distal TSS (TSS –2) was impacted to similar degrees as a stronger proximal TSS (TSS –1) by splicing perturbations of these SEs (Figure 5G). Together, these observations confirm that splicing of SEs can impact TSS use, particularly when the TSS is intrinsically weak, the SE is highly included, and the TSS is located proximal and upstream of the SE. The generalization of EMATS to SEs more broadly implies that gene expression may commonly be regulated through effects on splicing of promoter-proximal exons.

### Splicing factors impact TSS use and EMATs connection to neurogenesis

To investigate the potential biological role of regulation of gene expression via EMATS, we analyzed the functions of genes with EMATS structure. Interestingly, the 1777 mouse genes and the 1098 human genes with the strongest EMATS potential are enriched for functions in brain development, neuron projection, synapse organization and related functions (Figure 6A and S6A). This observation suggests that regulation via EMATS may contribute to neuronal differentiation, i.e. that splicing changes resulting from neuron-specific changes in splicing factor activity might trigger expression changes via EMATS that contribute to neuronal gene expression programs.

**Figure 6.**
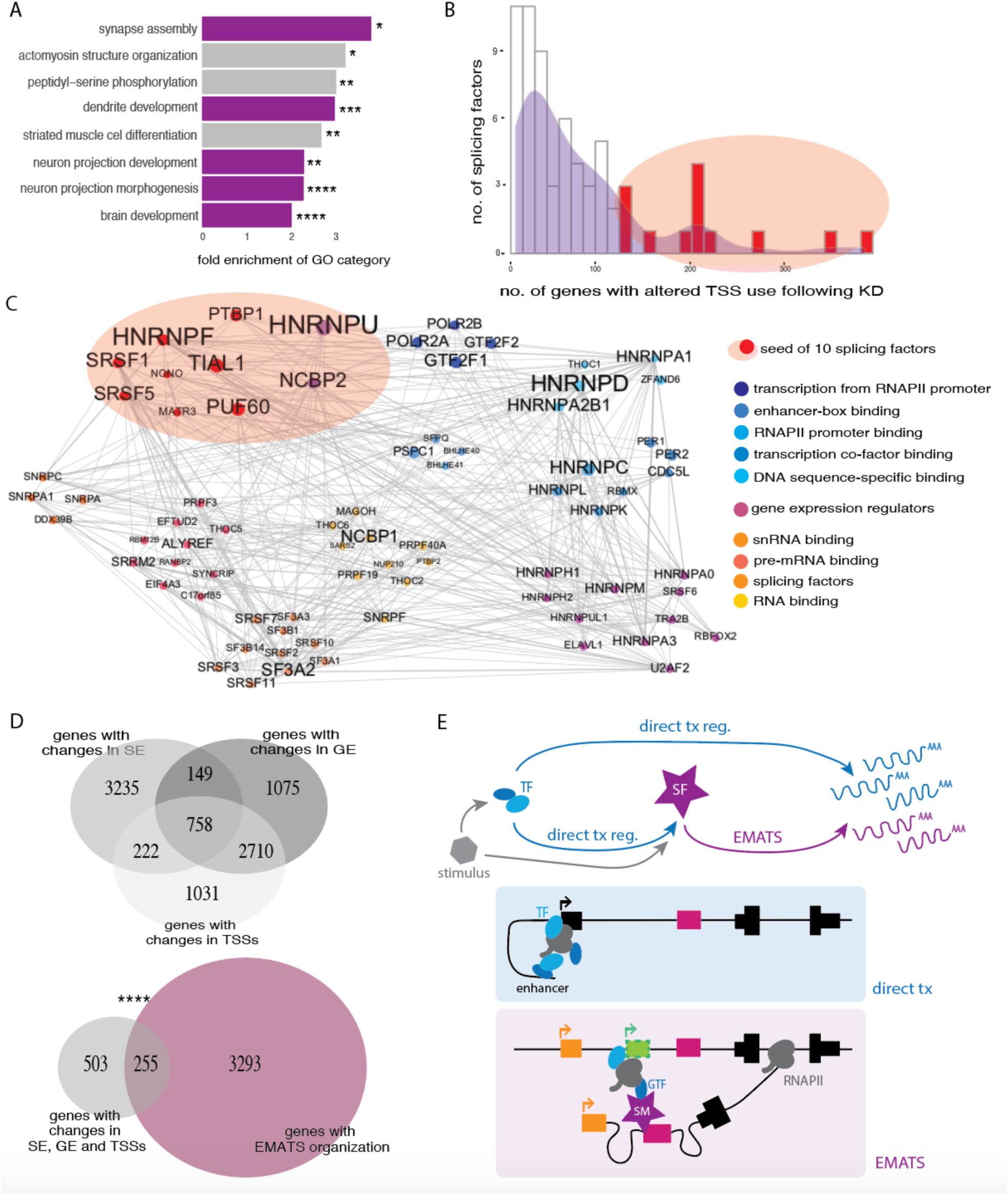
A subset of splicing factors impact TSS use and interact with transcription machinery. A, Gene Ontology analysis of 1777 mouse genes with the strongest EMATS potential. Fold enrichments shown for the most significant categories with asterisk indicating adjusted *p*-values and color indicating relation to neuron development. B, Histogram of number of genes with significant changes in alternative first exon usage following depletion of 67 splicing factors. Mean number between two cell lines (HepG2 and K562) is plotted for each RBP (top ten splicing factors with greatest number of changes shown in red). C, PPI network for the top 10 splicing factors from (B), colored by Gene Ontology category. Nodes represent proteins and edges represent PPIs. Node and label size are proportional to protein connectivity. The 10 selected splicing factors in red primarily interact with other 65 proteins, generating a network with 75 nodes and 424 edges, a diameter of 5, an average weighted degree of 6.1, an average clustering coefficient of 0.39 and an average path length of 2.1. PPI data are from STRING database (Szklarczyk et al., 2015) (using experimentally determined, database annotated, homology-based, gene fusion and automated text mining interactions). Networks were built using Gephi (http://gephi.org). D, (above) Venn diagram showing the overlap between genes with significant changes in gene expression (GE), alternative splicing of SEs and relative usage of TSSs following knockdown of *PTBP1* in human HepG2 cells. (below) Venn diagram showing the overlap between genes with changes in GE, SE and TSSs following knockdown of *PTBP1*, for human genes with EMATS organization. The overlap is 1.7-fold above background expectation (p < 1.6e-20). E, Model for the role of EMATS in dynamic gene expression programs. Growth factor or other stimuli activate transcription factors (TF) and splicing factors (SF). TFs influence gene expression by direct effects on transcription (tx) and indirectly by regulating levels of SFs. Effects of SFs on splicing contribute to gene expression programs by EMATS. In genes with EMATS structure, splicing machinery (SM) or SFs recruit GTFs or RNAPII to activate weak TSS(s) proximal and upstream of the exon. See also Figure S6.

The mechanistic link between splicing and TSS usage could be mediated by splicing machinery, splicing factors, or exon junction complex components, particularly those factors that interact with transcription machinery, transcription factors or chromatin. To explore potential links between splicing factors and TSS use, we analyzed transcriptome-wide changes in alternative TSS usage following knockdown of RNA-binding proteins (RBPs) using data from a recent ENCODE project (Van Nostrand et al., 2019). This analysis detected large numbers of TSS changes (Figure S6A), consistent with previous observations in *Drosophila* cells (Brooks et al., 2015). Depletion of factors involved in RNA splicing generally impacted larger numbers of TSSs than did depletion of other RBPs (Figure 6B and S6B). The ten splicing factors associated with the largest numbers of changes in TSS usage (Figure S6C) included PTBP1, whose downregulation plays a central role in neurogenesis (Linares et al., 2015). Using protein-protein interaction (PPI) data from the STRING database (Szklarczyk et al., 2015), we observed that these ten splicing factors interact with 65 other proteins, including subunits of RNAPII and GTFs (Figure 6C). Compared with the PPI partners of the ten splicing factors whose depletion affected the fewest TSSs, these 65 proteins were enriched for functions in enhancer binding, transcription factor activity and promoter proximal binding (Figure 6C, S6D and S6E). Together, these observations indicate that some splicing factors have wider impacts on promoter choice than previously recognized, and identify extensive interactions of these factors with core transcription machinery.

To investigate whether neuron-related splicing factors could regulate expression via EMATS, we analyzed transcriptome-wide changes following PTBP1 knockdown using the ENCODE data (Van Nostrand et al., 2019). Following PTBP1 knockdown, 758 genes had significant changes in SE splicing, TSS usage and gene expression, including 255 genes with EMATS organization; the latter represents a 1.7-fold enrichment over the background frequency of EMATS genes (Figure 6D). Among these 255 genes, the majority (165) also contained a PTBP1 eCLIP peak and had changes in TSS usage and gene expression that matched the direction expected from EMATS based on the direction of the change in splicing. For example, in the human *BMF* (Bcl2 Modifying Factor) gene we observed reduced exon inclusion accompanied by decreased use of upstream proximal TSSs and decreased gene expression following PTBP1 knockdown (Figure S6F). The results above suggest that splicing factors, including a key factor involved in neuronal differentiation, contribute to gene expression programs via EMATS regulation downstream of their effects on splicing.

## Discussion

Here, we have shown that creation of a new internal exon in a gene – during evolution, by directed mutation or by altered regulation – can activate transcription from an upstream TSS and thereby boost gene expression levels, a phenomenon which we refer to as EMATS. Our study highlights several features of this relationship: (i) it requires exon splicing, not merely presence of a 5’ or 3’ splice site; (ii) it is more potent when the exon is highly included and (iii) when the promoter is intrinsically weak; (iv) it is sensitive to genomic distance, occurring most robustly when exon and promoter are within 1-2 kilobases; and (v) the above features occur in thousands of mammalian genes (Table S3).

The most straightforward model to explain the above features (among other possible models) would involve direct positive effects of splicing components recruited to transcripts during transcription on recruitment of transcription machinery to nearby upstream promoters (Figure 6E). Splicing often occurs cotranscriptionally, and it is clear that splicing machinery and splicing factors are often recruited to nascent transcripts which are being transcribed and are therefore tethered to the gene locus. The splicing of exons can directly recruit core transcription machinery to the local vicinity, which may increase local concentration and occupancy of RNAPII at nearby promoters to increase transcription initiation (Damgaard et al., 2008; Fong and Zhou, 2001; Kwek et al., 2002). The involvement of splicing machinery or proteins deposited on the transcript in connection with splicing would explain feature (i) above, while the more efficient recruitment of splicing machinery to more efficiently spliced exons would explain feature (ii). Recruitment of RNAPII or GTFs might be expected to activate transcription more effectively at weaker promoters where RNAPII recruitment is limiting than at strong promoters with higher intrinsic RNAPII occupancy, explaining feature (iii). A requirement for direct physical interaction between splicing machinery and RNAPII or GTFs might constrain the genomic distances involved, feature (iv). However, the varied chromatin conformations of different gene loci – which, in some cases, might involve chromatin loops between promoters and alternative exons (Mercer et al., 2013; Ruiz-Velasco et al., 2017) – might alter distance requirements for different genes. Frequent occurrence of the evolutionary path outlined in Figure 4F and/or selection gene architectures that enable alternative 5’ UTRs (Singer et al., 2008), may explain the widespread occurrence of EMATS organization in mammalian genomes, feature (v).

The EMATS phenomenon has both evolutionary and regulatory implications. We propose that emergence of new internal exons and of new TSSs are linked (Figure 4F). Once so activated, the new TSS produces new transcript isoforms and higher overall expression of the gene in specific tissues, providing a substrate for the regulatory evolution of the gene. The most obvious regulatory role for EMATS would be as a means for splicing factors to contribute to gene expression programs involved in differentiation or cellular responses to stimuli (Figure 6E). Specifically, we propose that external stimuli such as growth factors or changes in cellular environment trigger gene expression changes not only via direct effects on TF activity (Malladi et al., 2016; Rajbhandari et al., 2018) but also by effects on splicing factor activity (Reinhardt et al., 2011; van der Houven van Oordt et al., 2000) or changes in splicing factor levels downstream of affected TFs, yielding additional gene expression changes via EMATS. An additional implication of our findings is that targeted activation (or repression) of the expression of a gene for research or therapeutic purposes may be achievable by use of compounds such as antisense oligonucleotides or small molecules (Havens and Hastings, 2016) that enhance or inhibit the splicing of an appropriately located promoter-proximal exon. Here, we have focused on EMATS involving alternative exons because of its endogenous regulatory potential, but such intervention could target appropriately positioned constitutive exons as well, roughly triplicating the number of potentially targetable genes. Recent studies have broadened the definition of enhancers, showing that some gene promoters also function as enhancers (Engreitz et al., 2016; Scruggs et al., 2015); our findings support further broadening of this definition to include some exons as well.

## Abbreviations used

RNAPII: RNA polymerase II
GTF: general transcription factor
CTD: carboxy-terminal domain (CTD) of RNAPII
snRNP: small nuclear ribonucleoprotein particles
PCPA: premature cleavage and polyadenylation site
TSS: transcription start site
EMATS: Exon-Mediated Activation of Transcription Starts
PSI: percent spliced in
Start-seq: high-resolution sequencing of polymerase-associated RNA
H3K4me3: histone 3 lysine 4 trimethyl
Kb: kilobases
MO: morpholino antisense oligonucleotides
SE: skipped exon
CE: constitutive exon

## Acknowledgments

We thank Alberto R. Kornblihtt, Phillip A. Sharp, and members of the Burge lab for helpful discussions and comments, as well as Maria Alexis, Jason Merkin, Peter Freese and Brenton R. Graveley for assistance with access to genomic data and analysis pipelines. This work was supported by grants from the NIH to C.B.B. (grant numbers HG002439, GM085319), and by the Pew Latin American Fellows Program in the Biomedical Sciences (A.F.).

## Author Contributions

A.F. and C.B.B designed the study and wrote the manuscript. A.F conducted computational analyses, designed and performed experiments. K.S.K. contributed technically to some experiments. C.B.B supervised the work.

## Declaration of Interests

The authors declare no competing interests. Correspondence and requests for materials should be addressed to

## STAR*METHODS

### CONTACT FOR REAGENT AND RESOURCE SHARING

Please direct any requests for further information and resources to the Lead Contact, Christopher B. Burge, Department of Biology, Massachusetts Institute of Technology, Cambridge MA 02138.

### EXPERIMENTAL MODEL AND SUBJECT DETAILS

#### Cell lines, cell culture and treatments

NIH3T3 and HeLa cells were grown in DMEM, with high glucose and pyruvate (Gibco), supplemented with 10% fetal bovine serum (FBS). Mouse CAD (Cath.-a-differentiated) cells were grown in DMEM/F12 (Gibco) supplemented with 10% FBS. For morpholino oligonucleotide (MO) treatment (Gene Tools), 20 µM of morpholino targeting 5’ or 3’ splice site or MO control was added with endoporter (Gene Tools) following manufacturer’s instructions to cells plated at low confluence and left for 24 h.

### METHOD DETAILS

#### RNA-seq analysis and genome builds

We used the RNA-seq data from 9 tissues from mouse and rat (3 individuals each) associated with Merkin et al. (Merkin et al., 2012), available at NCBI Gene Expession Omnibus (GEO) (accession no. GSE41637). Reads were mapped to the mm9 and rn4 genome builds, respectively, and processed using TopHat (Trapnell et al., 2009) and Cufflinks (Trapnell et al., 2012). Alternative splicing patterns and PSI values were analyzed using MISO (Katz et al., 2010). Exons were defined as in Merkin et al. (Merkin et al., 2012), requiring FPKM ≥ 2 and meeting splice site junction read requirements implicit in the TopHat mapping. Exons with 0.05 < PSI < 0.97 in at least one tissue and two individuals were categorized as skipped exons (SE). Exons with PSI > 0.97 in all expressed tissues were defined as constitutive exons (CE), if the gene was expressed in at least three tissues and two individuals. Genomic and splicing ages were defined as in Merkin et al. (Merkin et al., 2015) by the pattern of species with genomic regions aligned to the exon or with an expressed exon in the orthologous gene overlapping the aligned region, respectively, using the principle of evolutionary parsimony. Open reading frames (ORFs) were annotated as in Merkin et al. (Merkin et al., 2012) and used to classify exons as located in the 5’ UTR, 3’ UTR or coding region.

#### CRISPR sgRNA design, genetic deletions and genotyping

CRISPR-Cas cell lines with the 5’ splice site of *Stoml1* deleted were generated using the protocol described by Ran and coworkers (Ran et al., 2013). The single-guide RNA was designed in silico to target the 5’ splice site using the CRISPR Design Tool (http://tools.genome-engineering.org) and cloned into a Cas9 expression plasmid (pSpCas9). After transfecting CAD cells with the plasmid expressing Cas9 and the appropriate sgRNA, clonal cell lines were isolated and insertion/deletion mutations were detected by the Surveyor nuclease assay (IDT). Positive clones detected were amplified by PCR, subcloned into TOPO-TA plasmids, and individual colonies were sequenced to reveal the clonal genotype.

### RNA Extraction, RT-PCR and qPCR

Total RNA was extracted using the RNA-easy kit (Qiagen) according to the manufacturer’s protocol. Reverse transcription using M-MLV reverse transcriptase (Invitrogen) and random primers was performed according to the manufacturer’s instructions. For nascent RNA extraction, RNA was metabolically labeled with 5-Ethynil Uridine for 10 minutes using Click-iT (Invitrogen) and labeled RNA was extracted and amplified according to the manufacturer’s instructions. Quantitative PCR analyses were performed with SYBR green labeling using a LightCycler 480 II (Roche).

### ChIP and antibodies

Chromatin immunoprecipitation was performed using the MAGnify™ Chromatin Immunoprecipitation System (Invitrogen) according to the manufacturer’s recommendations. For each immunoprecipitation, we used 10 µg of H3K4me3 antibody (PA5-17420 from Invitrogen), 10 µg of RNA polymerase II antibody (Ab817 from Abcam), 10 µg of Transcription Factor IIF1 (GTFIIF1) antibody (PA5-30050 from Invitrogen) and 10 µg of Rabbit IgG antibody (Invitrogen) as a negative control. DNA was purified and quantitative PCR analysis was performed with SYBR green labeling using a LightCycler 480 II (Roche). Immunoprecipitated chromatin was normalized to input chromatin and control IgG antibody.

### 5’ RACE

5’ RACE experiments were performed with 5’ RACE System for Rapid Amplification of cDNA Ends (Invitrogen) using three gene-specific primers (GSP) that anneal to the known region and an adapter primer that targets the 5’ end. Products generated by 5’ RACE were subcloned into TOPO-TA vectors and individual colonies were sequenced.

### Plasmids and luciferase activity assay

Rat *Tsku* genomic region and mutants were cloned into the psiCHECK backbone. For transfection assays, 1 µg plasmid was transfected into each well of a 6-well culture plate using Lipofectamine 2000 (Life Technologies) according to the manufacturer’s recommendations and cells were harvested after 24 h. To measure luciferase activity, we used the Dual-Luciferase® Reporter Assay System (Promega).

### QUANTIFICATION AND STATISTICAL ANALYSIS

#### Definition of species-specific exons

Evolutionarily new exons were identified as is Merkin et al. (Merkin et al., 2015). Genomic mappings of mouse and rat RNA-seq data were combined with whole-genome alignments to classify the species distribution of exons. Only internal exons were considered in this analysis, excluding first and last exons, and only unique exons were considered, excluding exons that arose from intra-genic duplications to avoid issues related to possibly inaccurate genome assemblies, annotations or read mappings. In all, 1,089 mouse exons were classified as mouse-specific exons and 1,571 rat exons were classified as rat-specific exons, as they were detected in RNA-seq data from mouse or rat, respectively, but not from any other species analyzed (Supplementary Tables 1, 2). Most genes that contained a new exon had only one, with 159 mouse genes and 276 rat genes containing more than one new exon.

#### Transcription start site annotation

TSSs in mouse were identified using Start-seq data from Scruggs and coworkers (Scruggs et al., 2015) downloaded from GEO (accession no. GSE62151); Start-seq uses high-throughput sequencing of nascent capped RNA species from the 5’-end, allowing for definition of TSSs at nucleotide resolution. TSSs were defined in 2,000 bp search windows centered on RefSeq-annotated TSSs, using the location to which the largest number of Start-RNA reads aligned. Very closely spaced TSSs separated by less than 50 bp were considered as a single TSS in Figure 1D. To identify TSSs in the same RNA-seq data used to classify new exons, we used data from Merkin et al. (Merkin et al., 2012) (GEO accession no. GSE41637) mapped with TopHat combined with Ensembl annotations. As in Merkin et al. (Merkin et al., 2012), Cufflinks version 1.0.2 was used to identify novel transcripts. The set of TSSs from each library identified from transcripts as the start site of the first exon were combined with the existing Ensembl annotations and merged into a single set of annotations using Cuffcompare (Roberts et al., 2011). Cufflinks was then applied to each library to quantitate the same set of transcripts (Supplementary Table 3). The number of TSSs were also estimated by the number of H3K4me3 peaks assigned to each gene with ChIP data from Yu et al. (Yu et al., 2015) (GEO accession nos. GSE59896 and GSE59998).

#### Software for data analysis, graphical plots and statistical analyses

For data analysis we used R Bioconductor, BEDTools, SamTools, GenomicRanges and the Integrative Genomics Viewer. All statistical analyses were performed in R (v.3.4.2) and graphical plots were made using the R package ggplot2. Lower and upper hinges of box plots correspond to the 25^th^ and 75^th^ percentiles, respectively. The upper and lower whiskers extend from the hinge to the largest and lowest value no further than 1.5 × IQR (interquartile range), respectively. Notches give approximate 95% confidence interval for comparing the medians. Statistical significance is indicated by asterisks (*p < 0.05, **p < 0.01, ***p < 0.001, ****p < 0.0001, *****p < 0.00001).

### DATA AND SOFTWARE AVAILABILITY

#### Data availability

The RNA-seq data from 9 tissues from mouse and rat associated with Merkin et al. (Merkin et al., 2012) is available at GEO (accession no. GSE41637). The Start-seq data from Scruggs et al. (Scruggs et al., 2015) is available at GEO (accession no. GSE62151), as well as the H3K4me3 data from Yu et al. (Yu et al., 2015) (accession no. GSE59896 and GSE59998). PolyA-seq data from five mouse tissues is available in Derti et al (Derti et al., 2012). Data of evolutionarily new exons is available in Merkin et al. (Merkin et al., 2015) as well as here in Supplementary Tables 1 and 2.

### ADDITIONAL RESOURCES

#### New exon inclusion, TSS usage, and species-specific expression

We considered genes with new exons as all genes with a new exon with PSI > 0.05 in any of the 9 tissues sequenced. We grouped genes as control genes with no new exons and genes with new exons divided by whether the exon was included or excluded in a given tissue. We calculated the number of TSSs used in each gene in each tissue and considered genes that gained TSSs in mouse, genes that gained TSSs in rat, and genes with same number of TSSs in both species based on the numbers of TSSs for each species in each gene in each tissue, or when considering all tissues together. Gene expression in mouse was compared to that in rat by taking the ratio of expression in mouse to expression of the orthologous gene in the analogous tissue in rat.

### Definition of new exon-proximal cleavage and polyadenylation sites

Polyadenylation sites were identified using available polyA-seq data from five mouse tissues (brain, liver, kidney, muscle, testis) (Derti et al., 2012). Only reads aligning to unique loci were retained and ends of reads within 25 nt of each other on the same strand were clustered. Polyadenylation sites were considered to be new exon-proximal cleavage and polyadenylation (nePCPA) sites if they were located within 2 kb upstream or downstream of a new exon, and as skipped exon-proximal cleavage and polyadenylation (sePCPA) sites if they were located within 2 kb upstream or downstream of skipped exons.

### Effects on nascent and steady state RNA levels

Effects on transcription initiation should be reflected in nascent RNA, while effects on RNA stability would only be visible in steady state mRNA. In the *Tsku* gene, nascent RNA levels were reduced to a similar extent as steady state mRNA (Figure 2d, Extended data Figure 3b, Extended data Figure 5a-d), in both sense and antisense orientations. For other genes studied here, *Stoml1* and *Gper1*, we also observed similar effects on nascent RNA in sense and antisense directions (Figure 2c, Extended data Figure 3b, Extended data Figure 4a-c). Furthermore, the model invoking inhibition of PCPA involves U1 snRNP binding at a 5’ splice site, but we observed increased gene expression from creation of a 3’ splice site. Thus, our observations are consistent with splicing-dependent regulation of transcription initiation but not with models involving PCPA.

## KEY RESOURCES TABLE

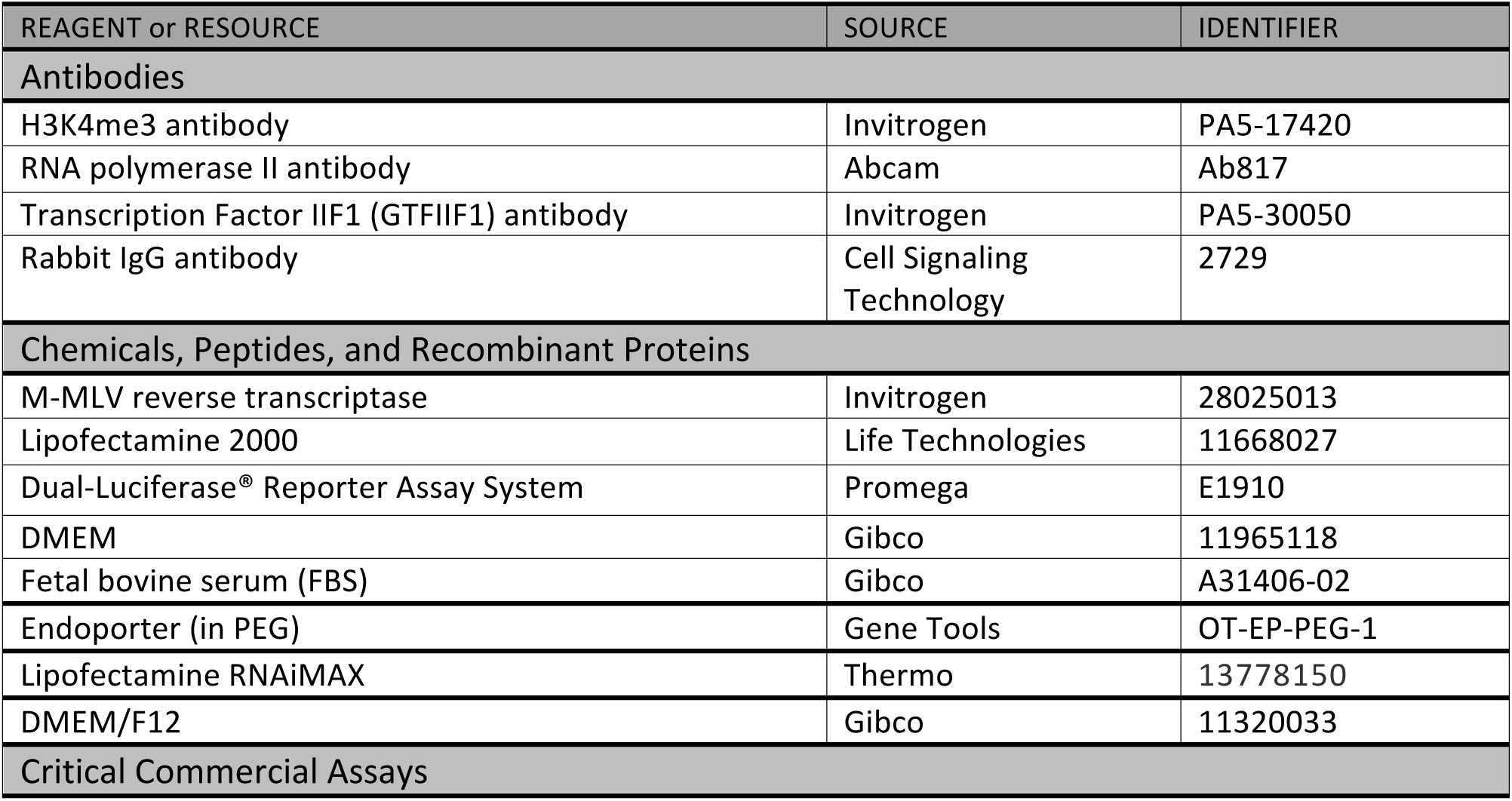
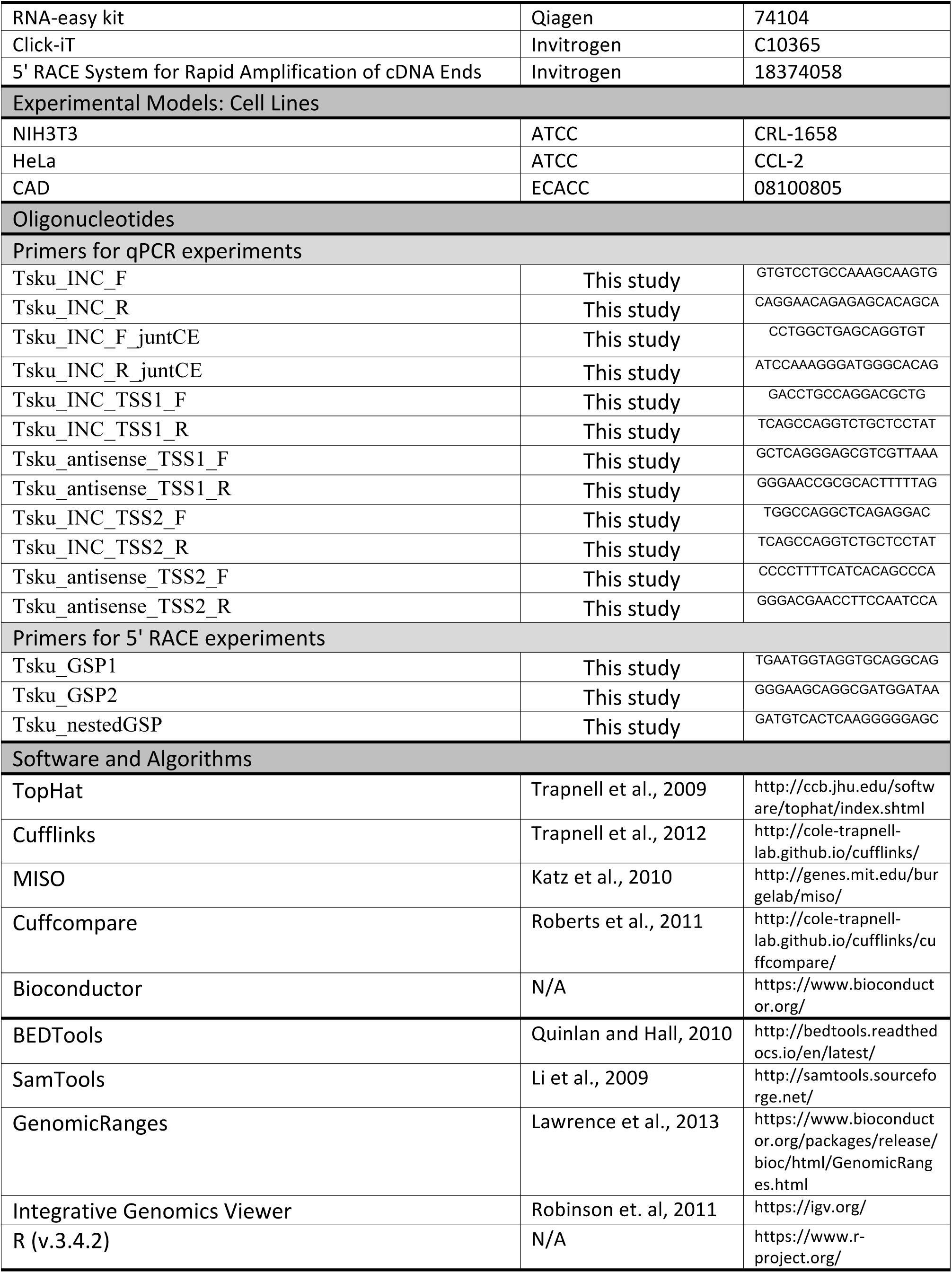

