## Supplemental Material for "Exon-mediated activation of transcription starts"

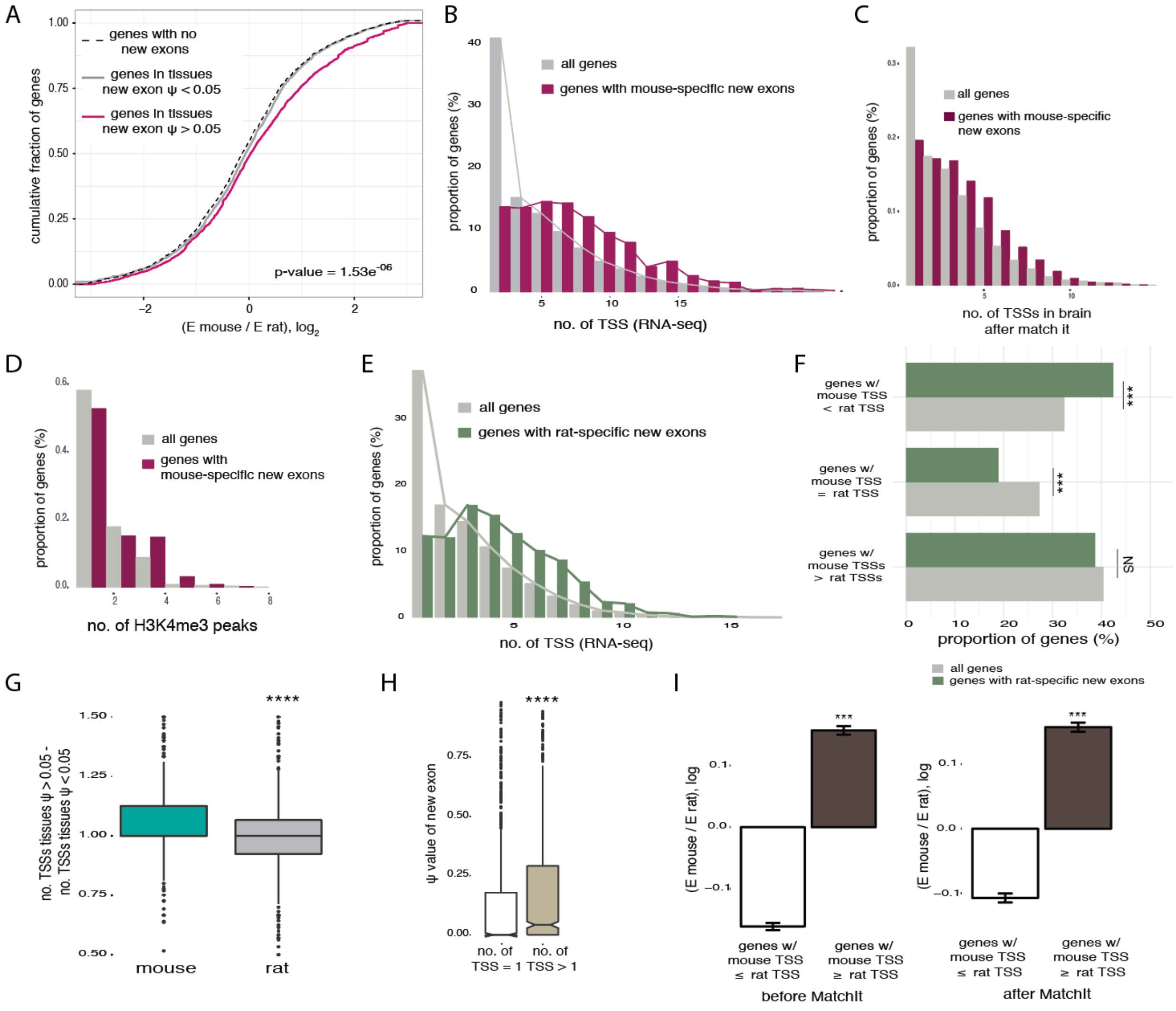


**Figure S1. Related to Figure 1**

A, Fold change in gene expression between mouse and rat for mouse control genes with no evolutionarily new exons (black, dotted line), genes with mouse-specific new exons in tissues where inclusion of the new exon is not detected, PSI < 0.05 (grey), and genes with new mouse-specific exons in tissues were the exon is included, PSI > 0.05 (pink). Statistical significance by Mann-Whitney U test is indicated between genes with mouse-specific new exons in tissues with PSI < 0.05 and tissues with PSI > 0.05. B, Distribution of the number of TSSs per gene using RNA-seq data across multiple species and multiple tissues, for all genes expressed in mouse and genes with mouse-specific new exons. Distributions are significantly different by Kolmogorov-Smirnov test. C, Distribution of the number of TSSs per gene in the mouse brain using RNA-seq data, for all genes expressed in mouse and genes with mouse-specific new exons, after matching the distribution of gene expression levels between the two groups using the MatchIt package in R. Distributions remain significantly different by Kolmogorov-Smirnov test after matching the gene expression levels between the groups, demonstrating that, independent of gene expression, genes with mouse-specific new exons are enriched in multiple TSSs. D, Distribution of the number of H3K4me3 peaks per gene using H3K4me3 ChIP-seq data for all genes expressed in mouse (grey) and genes with mouse-specific new exons (dark red). Distributions are significantly different by

Kolmogorov-Smirnov test. Genes with mouse-specific new exons are enriched in H3K4me3 peaks. E, Distribution of the number of TSSs per gene for all genes expressed in rat (grey) and genes with rat-specific new exons (green). Genes with rat-specific new exons are enriched in multiple TSSs (by Kolmogorov-Smirnov test). F, The proportion of genes with fewer TSSs in mouse (genes w/ mouse TSS < rat TSS), genes with the same number of TSSs in both species (genes we mouse TSSs = rat TSS), and genes that have more TSSs in mouse, for all genes expressed in both species (gray) and for genes with rat-specific new exons (green). Statistical significance indicated by asterisks corresponds to one-way ANOVA, Tukey post hoc test (NS = not significant). G, Fold change in the number of TSSs used per gene between tissues where mouse-specific exons are included (PSI > 0.05) and excluded (PSI < 0.05), for mouse genes and for the same tissues in rat. Evolutionary gain of internal exons and of TSSs are associated, but only in those tissues where new exons are included. H, Distribution of PSI values of new exons binned by the number of TSSs used in the same gene, for 9 tissues pooled together in mouse. I, Distribution of the fold change in gene expression levels between mouse and rat for genes with fewer or same number of TSSs used in mouse than rat (white) and genes with more TSSs used in mouse than rat (brown), before (left panel) and after (right panel) balancing the distribution of gene expression levels in mouse between the groups by MatchIt. Evolutionarily change in gene expression remain significantly different when balancing gene expression levels in mouse between groups, demonstrating that, independently of gene expression levels in one species, genes gaining TSSs in mouse have increased gene expression levels compared to rat.

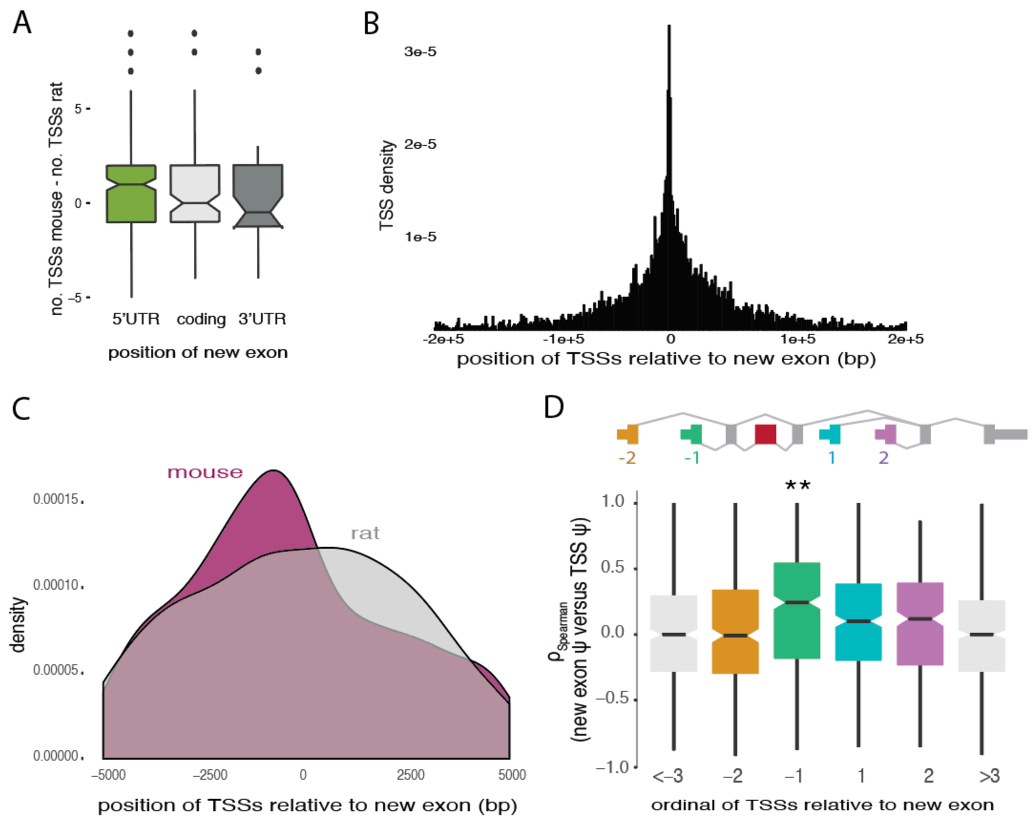


**Figure S2. Related to Figure 2**

A, Ratio between number of TSSs used in mouse and in rat for genes with mouse-specific evolutionarily new exons, binned by location of the exon within the gene. Increased number of TSS is associated with new exons located in the 5' UTR. B, TSS position relative to the start coordinate of the new exon in genes with mouse-specific new exons, for all TSSs used in 9 tissues in mouse. C, Comparison of distributions of TSS positions within 5 kb upstream and downstream of new exons between mouse (dark red) and rat (grey) for genes with mouse-specific new exons in all 9 tissues. The 0 position is at the start coordinate of the mouse-specific new exon (mouse) or at the location homologous to this position (rat). Distributions were smoothed with Kernel density estimation. D, Spearman correlations between the usage of a particular TSS and the PSI value of the new exon across multiple tissues for all TSSs used in genes with mouse-specific new exons, binned by their relative position to the new exon with negative numbers for TSSs located upstream of the new exon and positive numbers for TSSs located downstream of the new exon.

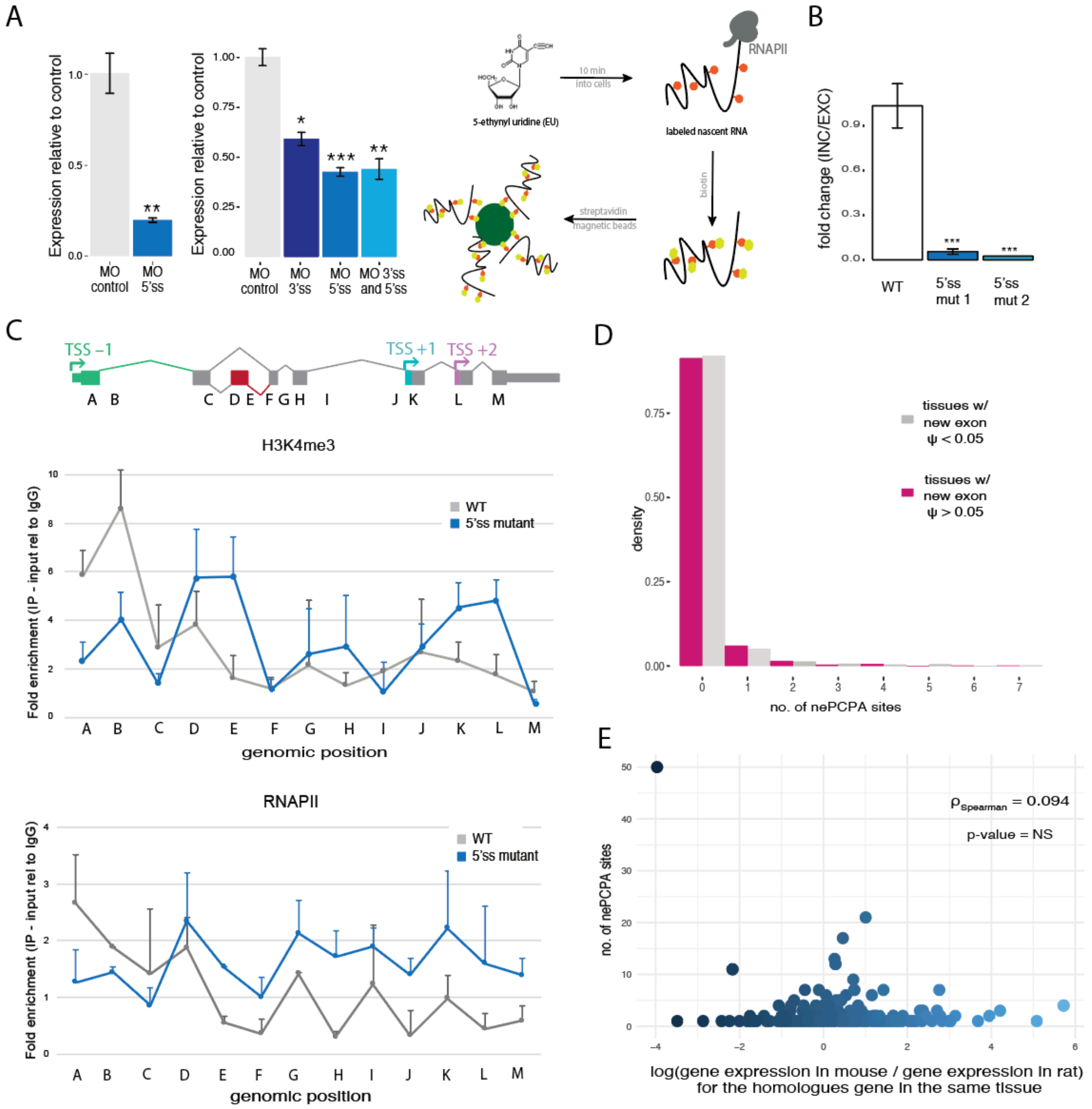


**Figure S3. Related to Figure 3**

A, A diagram representing the technique used to label nascent RNA with 5-ethynyl uridine and pull down the nascent RNA with the click-it method. Fold change in RNA levels of *Gpr30* (left) and *Tsku* (right) in NIH3T3 cells measured by qPCR. NIH3T3 cells were transfected with 20 µM morpholino (MO) targeting the 5' splice site of the new exon in Gpr30 or 20 µM MO targeting the 3' and/or the 5' splice sites of the new exon in *Tsku* for 24 h. Mean ± SEM of displayed distributions, *n*=3 biological replicates. Statistical significance indicated by asterisks corresponds to one-way ANOVA, Tukey post hoc test. B, Relative exon inclusion / exon exclusion of the mouse-specific new exon in *Stoml1* gene is shown, measured by qPCR of nascent RNA in wild type CAD cells and cells with CRISPR/cas-mediated mutations in the 5' splice site of the new exon. Mean ± SEM is shown for *n*=3 biological replicates. Statistical significance indicated by asterisks corresponds to one-way ANOVA, Tukey post hoc test. C, H3K4me3 and RNAPII profiles in Stoml1 gene in CAD cells determined by ChIP assay followed by qPCR with the regions indicated in the top panel. Values of two independent immunoprecipitations normalized to input and the mean value for control IgG antibody are shown for each region. Wild type cells (grey) and cells with CRISPR/cas-mediated mutations in the 5' splice site of the new exon (blue) are shown. D, Distribution of the number of polyadenylation sites used 2 kb upstream/downstream of new exons per gene in tissues new exon is excluded (PSI $< 0.05$, grey) and tissues with inclusion of new exons (PSI $> 0.05$, pink) for all genes with new exons. Distributions are not significantly different by Kolmogorov-Smirnov test. E, Scatter plot showing the relationship between the number of nePCPA sites and the fold change in gene expression levels between mouse and rat. These variables are not significantly associated by Spearman correlation test. Polyadenylation sites for 5 tissues in mouse were analyzed using polyA-seq data (Derti et al., 2012).

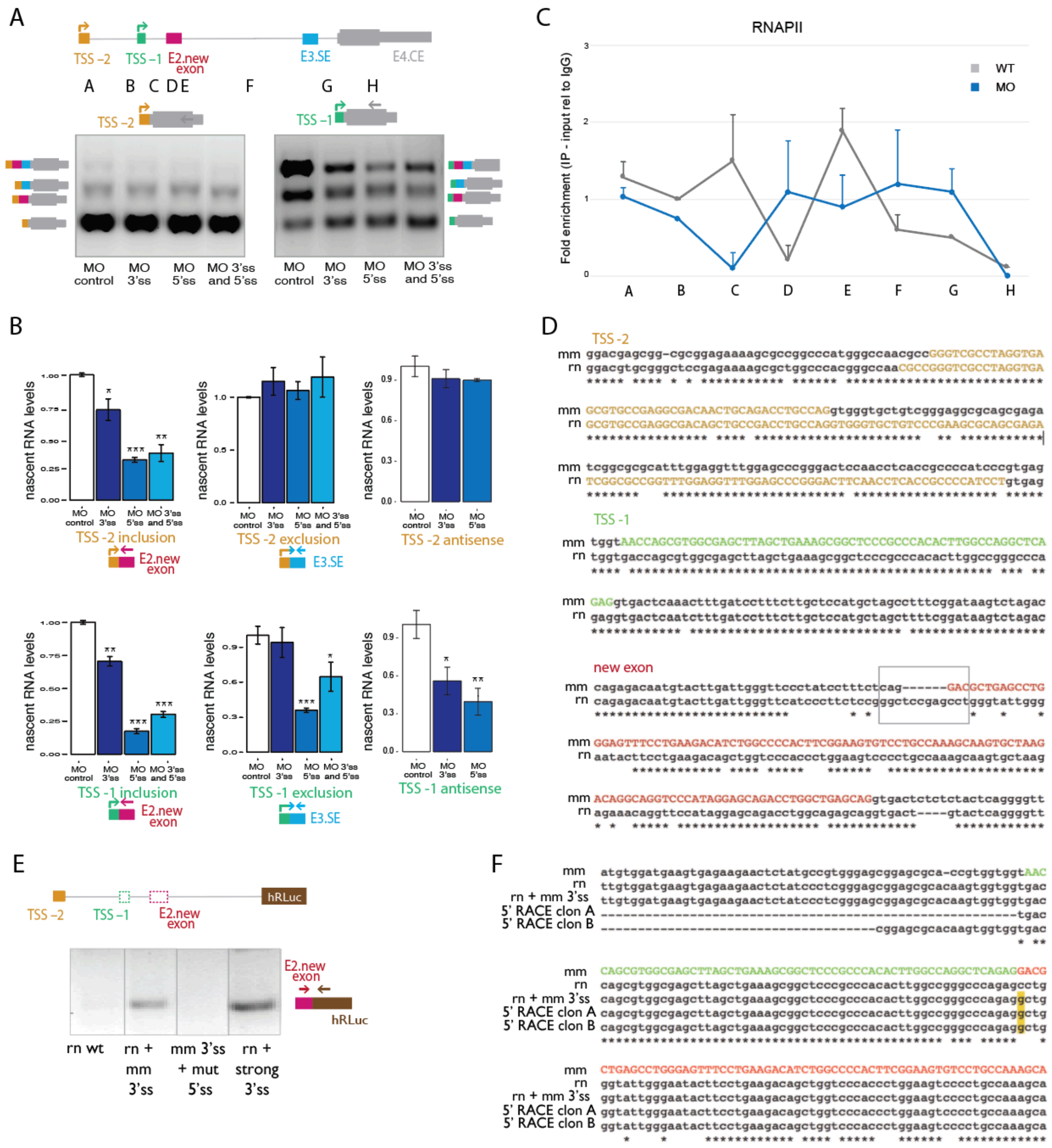

**Figure S4. Related to Figure 4**

A, Isoform expression for *Tsku* gene in NIH3T3 cells, measured by RT-PCR with primers targeting TSS –2 (top), TSS –1 (bottom) and exon E4.CE. Cells were transfected for 24 h with 20 μM MO targeting the 3' and/or 5' splice sites of the new exon as indicated. B, Fold change in inclusion and exclusion levels of the mouse-specific new exon in *Tsku* gene and antisense transcription levels from both TSSs measured by isoform-specific qPCR of nascent RNA with primers illustrated by arrows. Exon exclusion levels are measured from both alternative first exons to the following skipped exon downstream the mouse-specific new exon. NIH3T3 cells were transfected with control MO or MO targeting the 3' and/or 5' splice sites of the new exon. A decrease in the inclusion level of transcripts starting at TSS –2 is compensated by an increase in the exclusion levels, while total level of transcripts starting at TSS –1 is reduced by MO treatment. Mean ± SEM of displayed distributions, $n$=3 biological replicates. Statistical significance indicated by asterisks corresponds to one-way ANOVA, Tukey post hoc test. C, RNAPII profile in *Tsku* gene in NIH3T3 cells determined by ChIP assay followed by qPCR with the regions indicated in panel A. Values of two independent immunoprecipitations normalized to input and the mean value for control IgG antibody are shown for each region. NIH3T3 cells transfected for 24 hours with 20 μM control MO or MO targeting both 3' and 5' splice sites of the new exon. D, Alignments and identity between mouse (mm) and rat (rn) of the DNA sequence of TSS –2, TSS –1 and mouse-specific new exon in the *Tsku* gene. e, Splicing patterns of the *Tsku* gene in HeLa cells transfected with the hybrid constructs shown in Figure 4d. The creation of the mouse 3' splice site (rn + mm 3'ss) or of a stronger 3' splice site (rn + strong 3'ss) of the mouse-specific new exon in the rat sequence promotes the inclusion of the mouse-specific new exon in the rat context only when the wild type 5' splice site is maintained (but not in the mm 3'ss + mut 5'ss construct). F, Sequence of the 5' end of *Tsku* transcripts generated by 5' RACE in HeLa cells transfected with rat *Tsku* constructs with the 3' splice site of the mouse-specific new exon (5' RACE clone A, clone B) aligned to the mouse sequence (mm) and the rat sequence (rn). For 80% of the sequenced transcripts, the 5' end mapped 1 bp upstream of the position of mouse TSS –1 (clone A), while in the remainder the 5' end mapped 19 bp upstream of (clone B).

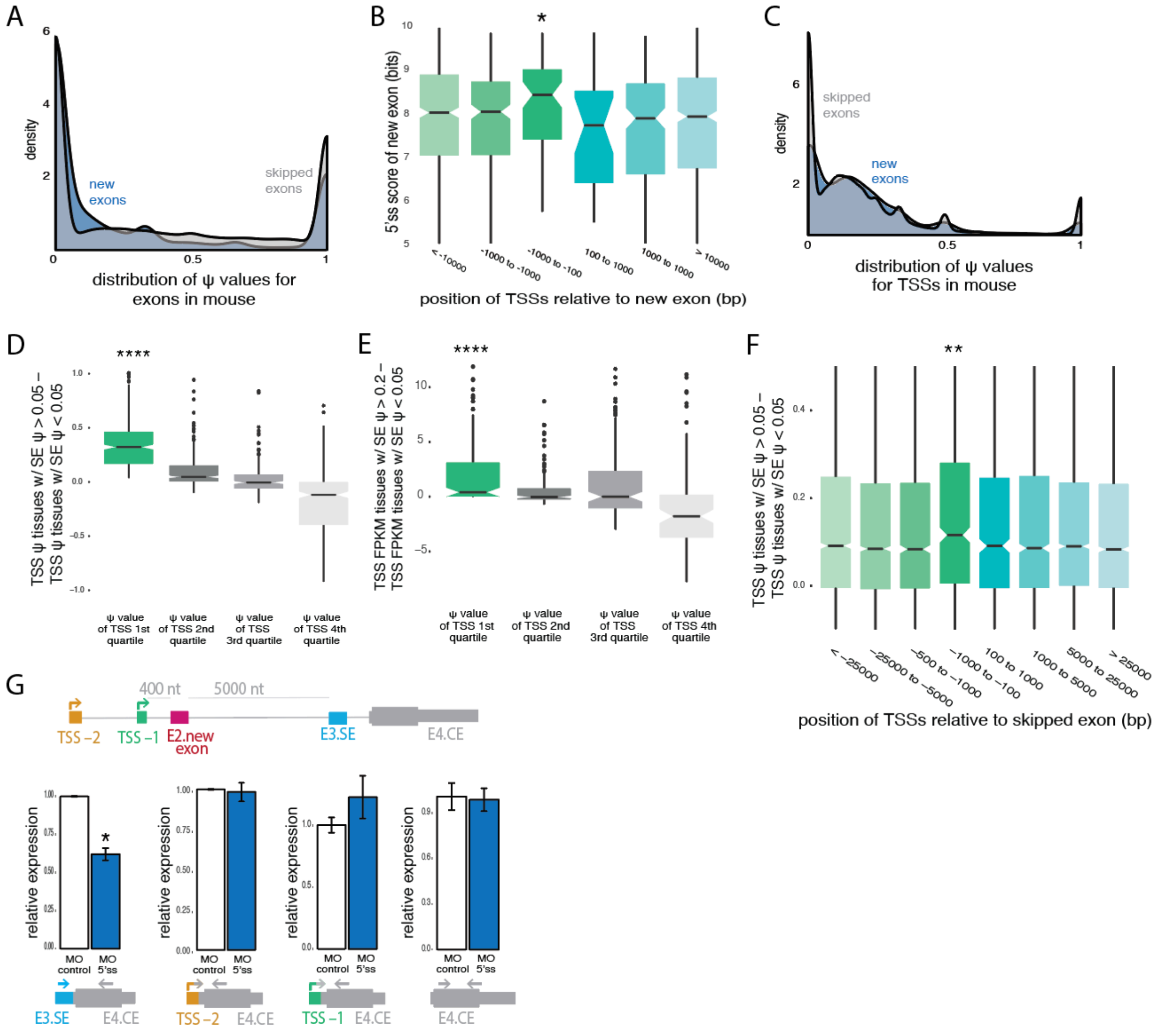


**Figure S5. Related to Figure 5**

A, Distribution of the PSI values of mouse-specific new exons (blue) and SE (grey) in mouse across 9 tissues. B, Distribution of 5' splice site scores of mouse-specific new exons, binned by the relative position to the next upstream TSS used in the same gene. 5' splice site scores were calculated using MaxEntScan (Yeo and Burge, 2004). C, Distribution of the PSI values of first exons associated with TSSs in genes with mouse-specific new exons (blue) and in genes with SEs in mouse (grey). D, E. Difference in TSS usage based on PSI value (d) and FPKM (e) in tissues with high versus low inclusion of skipped exons (SE), in the same gene across multiple tissues for proximal and upstream TSSs (within 1kb upstream the SE) used in genes with SEs in mouse, binned by quartiles of PSI values of the TSSs. F, Difference between TSS PSI values in tissues with high versus low inclusion of skipped exons (SE), for all weak TSSs (bottom quartile) used in genes with skipped exons in mouse, binned by their position relative to the SE. G, Fold change in inclusion (far left), exclusion levels (left center, right center) and total levels (far right) of the skipped exon in *Tsku* gene (E3.SE) from both TSSs measured by isoform-specific qPCRs with primers shown by arrows. Exclusion levels were measured from both alternative first exons relative to the next constitutive exon downstream of the skipped exon. NIH3T3 cells were transfected with control MO or MO targeting the 5' splice site of E3.SE. Mean ± SEM of displayed distributions, $n$=3 biological replicates. Statistical significance indicated by asterisks corresponds to one-way ANOVA, Tukey post hoc test.

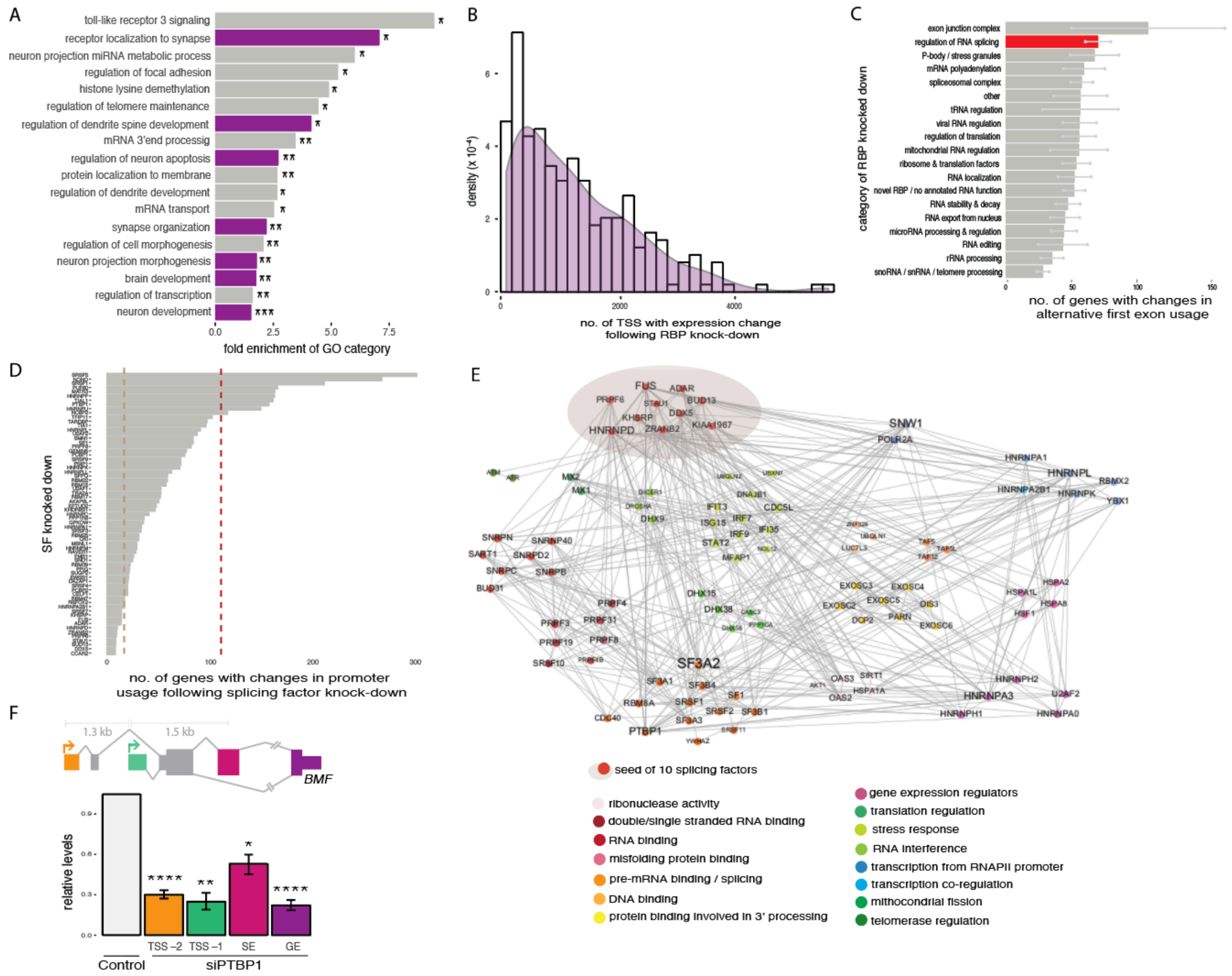


**Figure S6. Related to Figure 6**

A, Histogram and smoothed density of number of TSSs with significant expression change following depletion of each of 250 RNA binding protein genes. Mean two cell lines (HepG2 and K562) is plotted for each RBP. B, Distribution of the number of genes with significant changes in promoter usage associated with depletion of 250 RBPs, binned by Gene Ontology Biological Process categories of RBPs. Mean ± SEM between all RBPs in each GO category for two cell lines (HepG2 and K562) is plotted. C, Number of genes with significant difference in promoter usage associated with depletion of 67 splicing factors (SF). The red line indicates the cutoff for the top ten splicing factors driving the largest changes in promoter usage, while the green line indicates the cutoff for bottom ten control splicing factors driving the fewest changes in promoter usage. D, Protein interaction network for 10 control splicing factors driving the fewest changes in promoter usage. The control 10 splicing factors in red primarily interact with 88 other proteins, generating a network with 98 nodes and 410 edges, a diameter of 3, an average weighted degree of 4.29, an average clustering coefficient of 0.43 and an average path length of 1.32. Nodes represent proteins and links represent the interactions among them. Node size and label size is proportional to the protein connectivity (number of interactions a protein establishes with others). Protein interaction data were collected from STRING (Szklarczyk et al., 2015) and networks were built using Gephi (http://gephi.org). E, Gene Ontology analysis of 88 proteins included in the interaction network shown in (D), excluding the 10 seed control splicing factors. Adjusted p-values shown for the most significant categories with dotted line indicating adjusted *p*-value $< 0.05$. F, Exon-intron organization of human *BMF* gene. RNA-seq analysis of expression of *BMF* in HepG2 cells following PTBP1 knockdown normalized to expression of control cells. Inclusion levels of the skipped exon, as well as levels of relative usage of the alternative TSSs (TSS –2, TSS –1) and total gene expression are shown. Mean ± SEM of displayed distributions for $n=2$ replicates.

**Table S1. Related to Figure 1**
Mouse genes with new internal exons and their rat homologs. Columns A and H show the IDs for homologous mouse and rat genes with mouse-specific new exons, column B shows the locus of the new exon in mouse while column D shows the position of the new exon in the gene. Columns F and I show the average gene expression levels in brain for 3 individuals in mouse and rat, respectively. Column G shows the average PSI values in the mouse brain for 3 individuals.

**Table S2. Related to Figure 1**
Numbers of TSSs used in mouse and rat genes. Columns A and B show the IDs for homologous mouse and rat genes. Columns C and D show the numbers of TSSs used in mouse and rat, respectively, in a specific gene, pooling all nine sequenced tissues together.

**Table S3. Related to Figure 5**

| | mouse SE, TSS | SE ψ > median | TSS ψ < median | TSS located upstream | TSS < 2kb upstream |
|---|---|---|---|---|---|
| no. of SE | 49488 | 24744 | 13237 | 9621 | 3333 |
| no. of TSS | 58095 | 37266 | 18633 | 9510 | 2991 |
| no. of SE-TSS pairs | 223568 | 103801 | 42528 | 21326 | 4284 |
| no. of genes | 13491 | 9363 | 4973 | 3833 | 1777 |

Column A shows the number of SE expressed in the nine tissues sequenced in mouse and the number of genes in which they are distributed. Column B shows the number of these SE in which the average of PSI values across tissues is above the median of all SE and column C the TSS paired with those SE with an average PSI across tissues below the median. Columns D and E reflect the subset of SE-TSS pairs and genes from previous columns in which the TSS is located upstream or proximal and upstream of the SE.
